# White matter integrity requires continuous myelin synthesis at the inner tongue

**DOI:** 10.1101/2020.09.02.279612

**Authors:** Martin Meschkat, Anna M. Steyer, Marie-Theres Weil, Kathrin Kusch, Olaf Jahn, Lars Piepkorn, Paola Agüi-Gonzalez, Nhu Thi Ngoc Phan, Torben Ruhwedel, Boguslawa Sadowski, Silvio O. Rizzoli, Hauke B. Werner, Hannelore Ehrenreich, Klaus-Armin Nave, Wiebke Möbius

## Abstract

Myelin, the electrically insulating axonal sheath, is composed of lipids and proteins with exceptionally long lifetime. This raises the question how myelin function is affected by myelin turnover. We have studied the integrity of myelinated tracts after experimentally preventing the formation of new myelin in the CNS of adult mice, using an inducible *Mbp* null allele. Oligodendrocytes survived recombination, continued expressing myelin genes, but failed to maintain compacted myelin sheaths. Using 3D electron microscopy and mass spectrometry imaging we visualized myelin-like membranes that failed to incorporate adaxonally, most prominently at juxtaparanodes. Myelinoid body formation indicated degradation of existing myelin at the abaxonal side and at the inner tongue of the sheath. Compacted myelin thinning and shortening of internodes, with about 50% myelin lost after 20 weeks (=5 months), ultimately led to axonal pathology and neurological disease. These data reveal that functional axon-myelin units require the continuous incorporation of new myelin membranes.

## Introduction

Beyond the fundamental properties of white matter like increasing conduction velocity at low energy cost, myelin is much deeper involved in the organization and functioning of the brain and more dynamic than previously anticipated. Most of the myelin of the central nervous system (CNS) is formed by oligodendrocytes during early development (Emery, 2010). Apart from these early differentiated oligodendrocytes, new oligodendrocytes are continuously generated from persisting oligodendrocyte progenitor cells (OPCs) during adulthood (Crawford et al., 2014, Hughes et al., 2013, Young et al., 2013, Bergles and Richardson, 2015). The role of these adult formed oligodendrocytes in myelin plasticity during motor learning and the relevance for replacement of aged oligodendrocytes and their myelin is currently under intense investigation (Young et al., 2013, McKenzie et al., 2014, Bechler et al., 2017). Yet, since the early differentiated oligodendrocytes are long lived and survive alongside with the newly formed, it is less likely that oligodendrocytes differentiated in adulthood are mainly involved in the replacement and turnover of the developmentally formed myelin sheaths (Tripathi et al., 2017). Instead, they increase the total number of oligodendrocytes and add more myelin to the already existing white matter (Tripathi et al., 2017, Hill et al., 2018, Hill and Grutzendler, 2019), in agreement with a previous study in humans showing that a stable population of mature oligodendrocytes is established in childhood and remains static throughout life (Yeung et al., 2014). Notably the same study demonstrated that the myelin produced by this stable cell population is continuously exchanged and renewed. Since the myelin sheath is a large and tightly packed plasma membrane extension with limited accessibility, and since a single oligodendrocyte can maintain many myelin sheaths, it is plausible that its turnover occurs slowly. In a pulse-chase experiment using stable-isotope labeling with ^15^N and mass spectrometry long-lived proteins were identified in the rat brain (Toyama et al., 2013). Indeed, in contrast to cellular proteins that are turned over with a half-life of hours, myelin proteins such as PLP and MBP are long-lived and retain ^15^N in 20 % of the peptides after a chase period of 6 months. In a recent comprehensive study of protein lifetimes in the brain using in vivo isotopic labeling myelin proteins appeared in the extremely long lived protein population with a half-life between 55 days (2’,3’-cyclic-nucleotide 3’-phosphodiesterase (CNP)) and 133 days (claudin11 (Cldn11)) (Fornasiero et al., 2018). Targeted disruption of the *Plp*-gene in the adult showed that the abundance of PLP was halved within 6 months after tamoxifen-induction (Lüders et al., 2019). These data suggest that the individual myelin sheath is turned over and renewed by the respective oligodendrocyte in a continuous but very slow process.

Since the renewal of a myelin sheath is not well understood, we decided to directly visualize the turnover of myelin internodes in the adult mouse by ultrastructural analysis. To interfere with the maintenance of compact myelin, we generated a mouse line with a floxed exon 1 of the gene encoding myelin basic protein (MBP), which is common to all classical MBP isoforms (Takahashi et al., 1985, Pribyl et al., 1993, Campagnoni et al., 1993). MBP is an essential structural component of the CNS myelin by driving the adhesion of the cytosolic membrane leaflets that is required for the formation of multilayered compact myelin (Aggarwal et al., 2013, Raasakka et al., 2017, Vassall et al., 2015, Bakhti et al., 2014, Min et al., 2009). Upon compaction, MBP is thought to be the main constituent of the major dense line as observed by electron microscopy. Accordingly, the lack of MBP in the mouse mutant *shiverer* prevents myelin compaction (Readhead et al., 1987, Möbius et al., 2016, Rosenbluth, 1980). Therefore, oligodendrocytic processes only loosely associate with axons, but fail to establish a compact myelin sheath in a stable manner. Exploiting this as a structural distinguishing feature we obtained a tool to investigate the long-term stability and half-life of the individual compact myelin sheath. For this purpose, we crossed the MBP-flox line with the oligodendrocyte-specific inducible *Plp*-Cre^ERT2^ driver line (Leone et al., 2003). This inducible *Mbp* ablation allows us to eliminate MBP biosynthesis in mature oligodendrocytes to prevent the formation of novel compact myelin at an age when most developmental myelination has been achieved. After induction, MBP biosynthesis is abolished resulting in structural changes of the myelin sheath due to the lack of compaction of newly formed myelin membranes. We used mass spectrometry imaging of ^13^C-lysine pulse-fed mice by nanoscale secondary ion mass spectrometry (NanoSIMS), a technique to investigate the isotopic composition of the samples with high mass and lateral resolution (Agüi-Gonzalez et al., 2019) and found that the different myelin and axonal structures show different turnover rates. After *Mbp* ablation unusual enlarged inner tongue structures showed a higher content of ^13^C than compact myelin. In the current study, these structural transformations resembling a *shiverer*-like phenotype were investigated in detail by different electron microscopy techniques. With this approach we could reveal sites of insertion of newly synthetized myelin membranes, the manifestations of myelin removal and obtain an estimation of the half-life of a myelin internode in our model of adult myelin turnover. Determination and localization of myelin turnover reveals the dynamics and stability of this structure and helps to gain insight into the basic properties of the life of a myelin sheath during ageing.

## Results

### Adult MBP ablation induces a slow demyelination with survival of recombined oligodendrocytes

We generated mice that allow inactivation of the *Mbp*-gene in adult mice. For this purpose we established a mouse line with a lox-P flanked exon 1 of the classical *Mbp* locus (Mbp^fl/fl^) (Fig. 1A). By interbreeding with mice expressing tamoxifen-inducible Cre^ERT2^ in myelinating cells under the control of the *Plp* promotor (Leone et al., 2003) we gained control mice (Mbp^fl/fl^*Plp^*CreERT2wt*^) and inducible knockout mice (Mbp^fl/fl^*Plp^*CreERT2+*^), which were treated by i.p. injection of 1 mg tamoxifen per day at the age of 8 weeks for 10 days with a two days break in between (Fig. 1B). Genomic PCR analysis of brain lysate at 6 months after tamoxifen-induction confirmed that recombination of the floxed allele took place only in (Mbp^fl/fl^*Plp^*CreERT2+*^) mice (Fig. 1C), termed hereafter inducible conditional knockout mice (iKO) in comparison to tamoxifen-injected (Mbp^fl/fl^*Plp^*CreERT2wt*^) control mice. Time points of analysis after induction are indicated as weeks post tamoxifen induction (pti).

**Figure 1:**
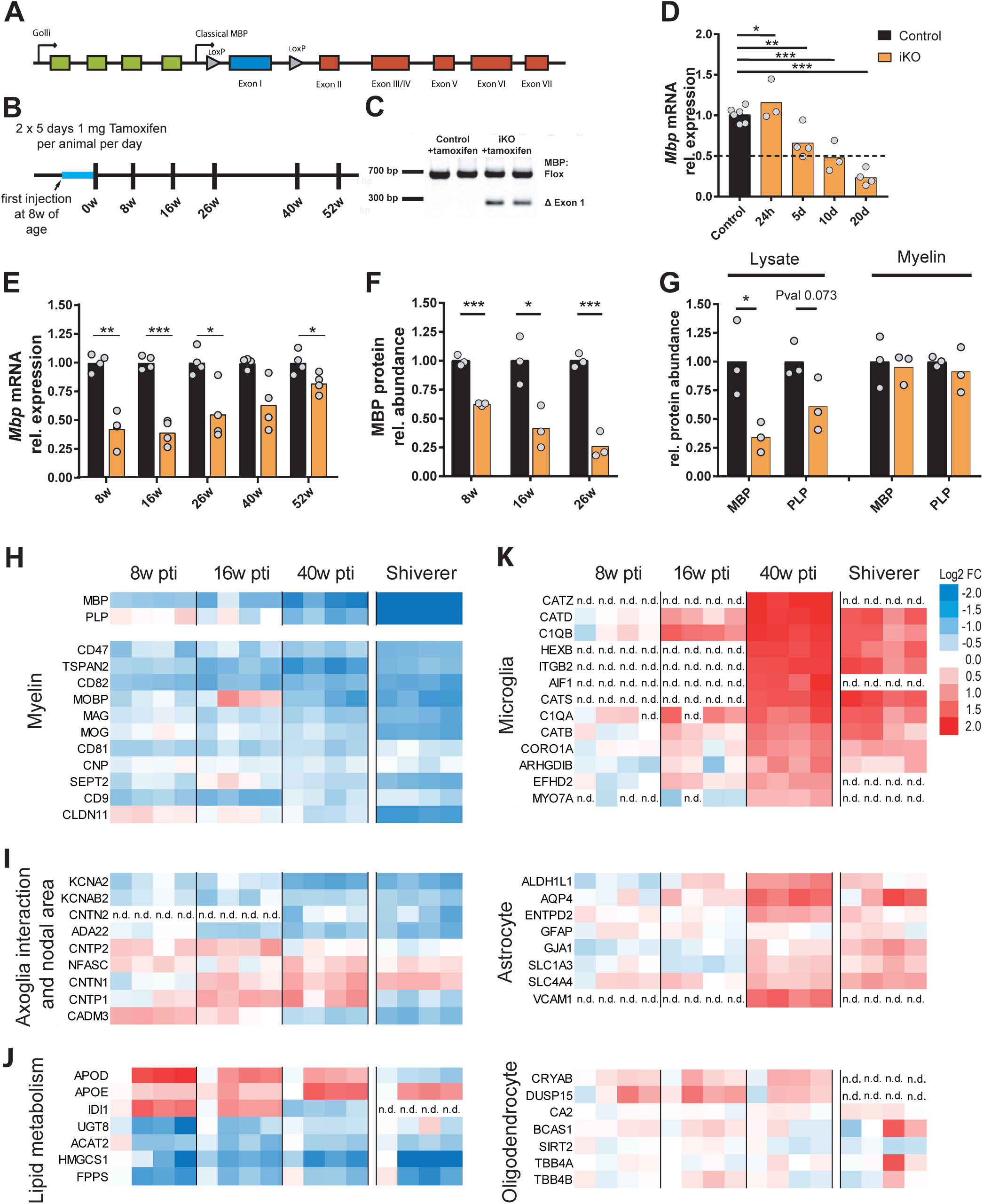
Deletion of the *Mbp* gene in mature oligodendrocytes and subsequent molecular changes. **(A)** Schematic of *Mbp* gene structure with floxed exon 1. **(B)** Experimental design and time points of analysis **(C)** Floxed exon 1 of the classical *Mbp* locus is deleted upon tamoxifen injection using the inducible *Plp*-CreERT2 mouse line (Leone et al., 2003, *Mol Cell Neurosci* 22:430-440). **(D)** and **(E)** Relative *Mbp* mRNA abundance in brain lysate at the indicated time points post tamoxifen injection (pti). The stippled line indicates reduction to 50%. **(F)** Relative abundance of MBP in brain lysate at indicated time points by immunoblot analysis **(G)** Immunoblot analysis of MBP and PLP abundance in lysate and myelin fraction 26 weeks pti. The protein abundance in the myelin fraction is unchanged (see also Figure S1 and 2) (Two-tailed unpaired t-test, p<0.05 (*), p<0.01 (**), p<0.001 (***)). **(H-K)** Heatmaps of normalized abundance of proteins selected from the quantitative proteome analysis of whole optic nerve in iKO at indicated time points and *shiverer* mice at the age of 10 weeks. Myelin proteins **(H)**, proteins involved in axoglia interaction and present in the node area **(I)** and proteins of lipid metabolism **(J)** are depicted. Markers of microglia, astrocytes and oligodendrocytes **(K)** were assigned to the cell type according to Zhang et al., 2014. Shown are the averages of two technical replicates from N=4 mice, optic nerve lysate, 8, 16 and 40 weeks pti and *shiverer* at 10 weeks of age. For normalization, iKO abundance values were divided by the mean of the corresponding control group with the color code representing down-(blue) or up-regulation (red) as log_2_-transformed fold-change. Abundance values for MBP and PLP were derived from a dataset recorded in the MS^E^ acquisition mode dedicated to correct quantification of exceptionally abundant proteins (see also Fig S3, S4 and S5, Supplementary table 1 and STAR Methods for details), n.d. not detected.

The relative abundance of *Mbp* mRNA was determined in the brain 24 h and 5, 10 and 20 days after the first tamoxifen injection (Fig. 1D). The mRNA level declined to 50% after 10 days and 23 % of the control expression level within 20 days in the brain. As the mice aged, the *Mbp* mRNA abundance in the brain partly recovered to approx. 40% of the control at the time points 8 and 16 weeks pti. Later, mRNA levels increased further to 55% at 26 weeks, 63% at 40 weeks and 82% at the latest time point of 52 weeks pti, suggesting *Mbp* expression by oligodendrocytes differentiated in adult iKO mice from non-recombined oligodendrocyte precursors (OPC) (Fig.1E). The expression levels of the mRNA encoding other myelin proteins PLP, MAG, MOG and CNP were largely unchanged (Fig. S1A). The ablation of exon 1 of the classical *Mbp* gene did not impair the expression of *Golli* mRNA which is encoded by the same transcription unit (Fig. 1A and Fig. S1A).

Since myelin proteins are known to exhibit a long life time, we next analyzed the MBP protein abundance in total brain lysate in our mouse model of adult *Mbp* ablation. As expected, MBP levels progressively decreased showing a significant reduction at 8, 16 and 26 weeks pti (Fig. 1F). From this immunoblot analysis, we calculated a half-life of about 77 days (11 weeks). Other myelin proteins PLP, MAG and MOG also decreased in abundance in brain lysate, but to a lesser extent (Fig. S1B-D). 26 weeks pti, MBP levels were reduced to 26 % of control. At this time point, the iKO mice began to show a progressive motor phenotype characterized by tremor and ataxia (data not shown).

To discriminate between the possibilities that MBP is either lost from the myelin sheath or that the amount of compact myelin itself is decreased, we purified myelin from brain lysate and analyzed the relative abundance of MBP and PLP. As shown in Fig. 1G and Fig. S2A 26 weeks pti MBP abundance was significantly reduced in brain lysate. In contrast, MBP levels were not changed in a purified myelin fraction. Moreover, compared to controls, the myelin fraction of iKO mice was visibly reduced, indicating a loss of compact myelin (Fig. S2B and C).

To determine whether this *Mbp*-ablation induced demyelination was caused by the loss of oligodendrocytes, we investigated oligodendrocyte numbers and the proliferation of OPCs. The determination of OPC and oligodendrocyte numbers by labeling Olig2 and PDGFRA, respectively (Fig. S3A and B) was performed in the fimbria, a comparatively homogenous white matter tract in the brain 46 weeks pti. We found a significantly increased density of Olig2^+^ and PDGFRA^+^ cells in the iKO while the area of the fimbria remained unchanged (Fig. S3C). In addition, significantly increased numbers of CAII positive oligodendrocytes in the iKO were found 26, 46 and 52 weeks pti (Fig. S3D). In accordance, TUNEL staining did not indicate increased apoptosis of cells at 40 and 52W pti (Fig. S4A and B). To track proliferating cells we administered 5-ethynyl-2’-deoxyuridine (EdU, 0.2 mg/ml in drinking water) at the time point 40 weeks pti for 3 weeks followed by a EdU-free chase period of 3 weeks. Double staining with EdU revealed that the percentage of EdU-positive Olig2^+^ and PDGFRA^+^ cells increased 4-fold in the iKO (Fig. S3E, E’ and F, F’). Approximately 9% of the PDGFRA^+^ cells were also EdU positive in the iKO compared to 2.7 % in the control, (Fig. S3F and F’). However, within the period of the EdU administration and chase these OPCs did not differentiate to CAII positive oligodendrocyte in significant numbers (Fig. S3G and G’). Expression analysis in the corpus callosum 40 and 52 weeks pti showed unchanged or increased expression of PLP, Olig2, PDGFRA and CAII, while MBP expression was significantly reduced (Fig. S4C). We conclude that recombined oligodendrocytes persist and continue cell type specific protein expression while newly differentiated oligodendrocytes might partially account for an increase in myelin gene transcripts.

### Whole optic nerve proteome analysis reveals similarities between *Mbp* iKO and *shiverer* mice

To obtain information about changes in the abundance of myelin proteins and about the systemic response to this slow demyelination by adult *Mbp* ablation in a more systematic way, we utilized quantitative mass spectrometric proteome analysis. We chose the optic nerve as a suitable model CNS white matter tract because of its high degree of myelination and the possibility to extract the complete intact structure. Moreover, different to corpus callosum the myelination pattern remains stable once the developmental myelination is completed. Therefore, it served as model tissue for myelin maintenance also in the subsequent fine structural investigation since it can be prepared for electron microscopy analysis by chemical fixation as well as high-pressure freezing allowing for a wide range of ultrastructural analyses.

For proteome analysis, we obtained whole optic nerve lysates from iKO and controls at 8, 16 and 40 weeks pti and for comparison also optic nerve lysates from 10 weeks old *shiverer* mice, which suffer from inability to developmentally form regular compact myelin due to the lack MBP (Rosenbluth, 1980) (Fig. 1H-K). At this time point, *shiverer* mice reach the clinical end stage. In total, we identified and quantified 1863 proteins with an average sequence coverage of 38.5% from the iKO/control samples of the three time points, and 1690 proteins with an average sequence coverage of 33.8% from the *shivere*r/control samples (Supplementary Table 1 and Figure S5).The proteins were assigned to the enriched expression in cell types according to the RNA-Seq transcriptome (Zhang et al., 2014). Apart from analyzing the protein abundance changes upon MBP deletion for each time point individually (iKO vs Ctrl; Supplementary Table 1), we also compared early (8 weeks) and late (40 weeks) iKO/Ctrl ratios to detect differences in normalized protein abundance over the course of MBP deficiency (iKO/Ctrl 40w vs iKO/Ctrl 8w; Figure S5, Supplementary Table 1). Guided by this analysis, we selected proteins of interest from the entire iKO/Ctrl dataset and compared their normalized abundance with that in *shiverer* mice as a proxy for the demyelination endpoint (see heatmaps in Fig. 1). Indeed, we found numerous myelin proteins reduced in abundance (MBP, PLP, MAG, MOG, CNP, claudin 11 (CLDN11), CD9 and tetraspanin-2 (TSPAN2) (Fig. 1H), while markers of oligodendrocyte cell bodies were unchanged or slightly elevated (carbonic anhydrase 2 (CAH2), BCAS1, CRYAB) (Fig. 1K). Indicative of neuropathology, levels of microglial markers (cathepsins, iba1 (AIF1), HexB, complement subcomponents C1QB and C1QA)) and astrocyte markers (ALDH1L1, AQP4) were increased at the late time point in the iKO (Fig. 1K). Virtually complete absence of MBP was confirmed in the *shiverer* optic nerve proteome (Supplementary Table 1). In addition, we found a strong reduction in the amount of other myelin proteins also in *shiverer* such as PLP, claudin11, septin2, septin4, MAG and tetraspanin-2. The almost unchanged presence of oligodendrocyte markers SIRT2 and carbonic anhydrase 2 confirmed that MBP expression is not required for oligodendrocyte survival (Fig. 1K). The abundance of microglial enriched markers was similarly elevated in *shiverer* as 40 weeks pti in iKO mice. In addition, astrocytic markers were increased in *shiverer* indicating astrogliosis. Interestingly, chronic MBP deficiency and induced loss of MBP 40 weeks pti showed similarities in the proteome with elevated levels of proteins involved in axo-glia interaction at the paranode like contactin-1 (CNTN1) and neurofascin (NFASC) and decreased amounts of CADM3 at the internode (Fig. 1I). In the iKO as well as in *shiverer* also the levels of juxtaparanodal voltage-gated potassium channel α subunit K_v_1.2 (KCNA2) and the subunit K_v_β2 (KCNAB2) were decreased. Furthermore, we found a significantly diminished abundance for their interaction partner ADAM22 and for contactin-2 (CNTN2), indicating alterations in the juxtaparanodal organization.

As depicted in Fig.1J we also detected abundance changes in proteins involved in lipid metabolism. These comprise enzymes of the isoprenoid and cholesterol biosynthetic pathway such as HMG-CoA-synthase (HMGCS1), isopentenyl-diphosphate delta-isomerase 1 (IDI1) and farnesyl pyrophosphate synthase (FDPS). Abundance of HMG-CoA-synthase 1, which catalyzes the rate-limiting step in this pathway, was reduced in iKO as well as in *shiverer*. In addition, levels of the apolipoproteins Apo D (APOD, expressed in oligodendrocytes) and Apo E (APOE, expressed in microglia and astrocytes), both involved in cholesterol transport, were found increased in the iKO. Another enzyme important for myelin lipid synthesis, UDP-galactose:ceramide galactosyl-transferase (UGT8), was reduced in abundance. Taken together, after ablation of MBP the subsequent loss of myelin proteins in this mouse model was accompanied by changes in proteins involved in axo-myelinic interaction and myelin lipid synthesis. Similarities in the whole optic nerve proteome of *shiverer* and iKO mice suggest that progressive demyelination by the deletion of MBP in adulthood induced a state that resembles the dysmyelinated situation in *shiverer* in many aspects. However, the *Mbp* iKO mouse allows the morphological assessment of demyelination.

To validate the indications of neuropathology found by proteome analysis we investigated neuropathology by immunohistochemistry of GFAP, MAC3, APP and CD3 in the fimbria (Fig. S6). An increase in the GFAP-immunopositive area was detected 16 weeks pti together with a significant increase in the number of CD3 immunopositive cells, followed by increased MAC3-immunopositive area and the appearance of APP-positive spheroids 26 weeks pti. These signs of neuropathology were progressive. In conclusion, the ablation of MBP in mature oligodendrocytes caused a slowly progressing demyelination without impairment of myelin gene expression or oligodendrocyte survival, accompanied by a slow development of neuropathology, validating our mouse model as suitable for a fine structural analysis of the maintenance of the myelin sheath.

### Determination of turnover by metabolic stable-isotope labeling of iKO mice and NanoSIMS

Next we used the *Mbp* iKO model to directly visualize the integration of newly synthetized proteins into the mature myelin sheath using NanoSIMS. For this purpose we applied stable isotope labeling by pulse-labeling iKO mice for 45 days with a ^13^C-lysine diet starting at the age of 28 weeks (18 weeks pti) according to (Fornasiero et al., 2018) followed by one week of chase with normal diet and collected tissue at 26 weeks pti. Because of the limited lateral resolution of NanoSIMS we used spinal cord samples which contain large myelinated fibers. The samples were prepared for transmission electron microscopy and mapped for the occurrence of phenotypical changes of the compact myelin structure due to the lack of MBP. As the most striking difference to the control we found a tubular-vesicular enlargement of the inner tongue in the iKO sample (Fig 2A). The EM images were correlated with images of the ^12^C and ^13^C distribution. By selecting regions of interest the local ratio of ^13^C to ^12^C was determined on several morphological categories as described in Fig. 2. In detail, we assessed the ^13^C to ^12^C ratio on structures in the enlarged inner tongue (Fig. 2A and C), on compact myelin, myelin debris and a myelinoid body (Fig. 2B and C). Compact myelin structures show less enrichment of ^13^C than the axon and the structures in the enlarged inner tongue (Fig. 2C). This indicates that proteins in the axon and the inner tongue structures were turned over faster compared to the compact myelin sheath.

**Figure 2:**
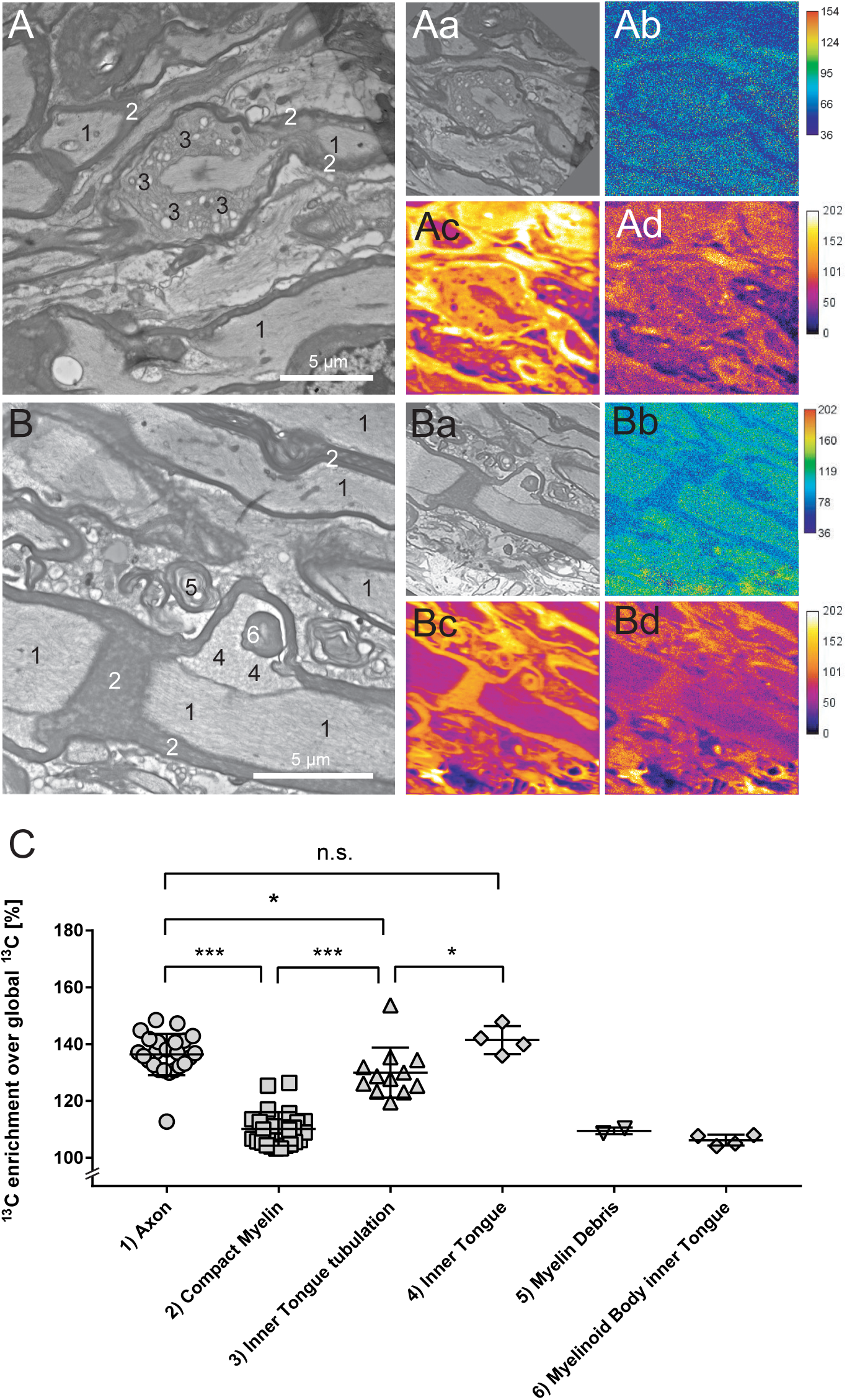
Determination of myelin turnover by ^13^C-lysine feeding and NanoSIMS imaging in *Mbp* iKO. Longitudinal spinal cord TEM section of iKO fed for 45 d with ^13^C-Lys SILAC diet and sacrificed after 1 week of chase with non-labeled control diet at 26 weeks pti. At the indicated regions of interest small areas were sampled and the enrichment of ^13^C was calculated from the NanoSIMS images. **(A)** Tubular-vesicular enlargement of the inner tongue. **(B)** A myelinoid body is visible at the inner tongue. **(Aa and Ba)** Aligned TEM image **(Ab and Bb)** ratio of ^13^C/^12^C **(Ac and Bc)** ^12^C NanoSIMS image **(Ad and Bd)** ^13^C NanoSIMS image. **(C)** ^13^C enrichment of the sampled areas. Every data point corresponds to a sampled area drawn on the TEM image (two*-*tailed unpaired t*-*test (C) p < .05 (*), p < .01 (**), p < .001 (***)). Scale bars: (A and B) 5 μm

### Compact myelin diminished by insertion of non-compacted membranes, myelin sheath thinning and internode shortening

To investigate the conspicuous structures at the inner tongue in detail, we assessed this phenotype in iKO mice by ultrastructural analysis of the optic nerve. In high-pressure frozen samples, we observed demyelination at 16 weeks pti that became widespread at 26 weeks pti (Fig. 3A). Alongside with demyelinated axons the emergence of membranes processes resembling the *shiverer* phenotype was observed that appeared most obvious at the myelin inner tongue or both at inner and outer tongue and looked alike the structures detected by the NanoSIMS analysis in the spinal cord. Only occasionally membrane tubules could also be found adjacent to non-myelinated axons (Fig. 3B and C). Such *shiverer*-like membrane tubules often left residual myelin sheaths behind that incompletely covered the axon (lower left panel in Fig. 3B). We quantified the occurrence of myelin tubulations at 16 weeks pti and assigned a location, i.e. the inner tongue, inner and outer tongue and adjacent to axons (Fig. 3C). Outer tongue tubulations were only clearly identified where inner and outer tongue tubulations occurred at the same myelinated axon (Fig. 3B, upper right panel). Tubules found in close vicinity to non-myelinated axons were counted as “adjacent to axons” (Fig. 3B lower right panel).

**Figure 3:**
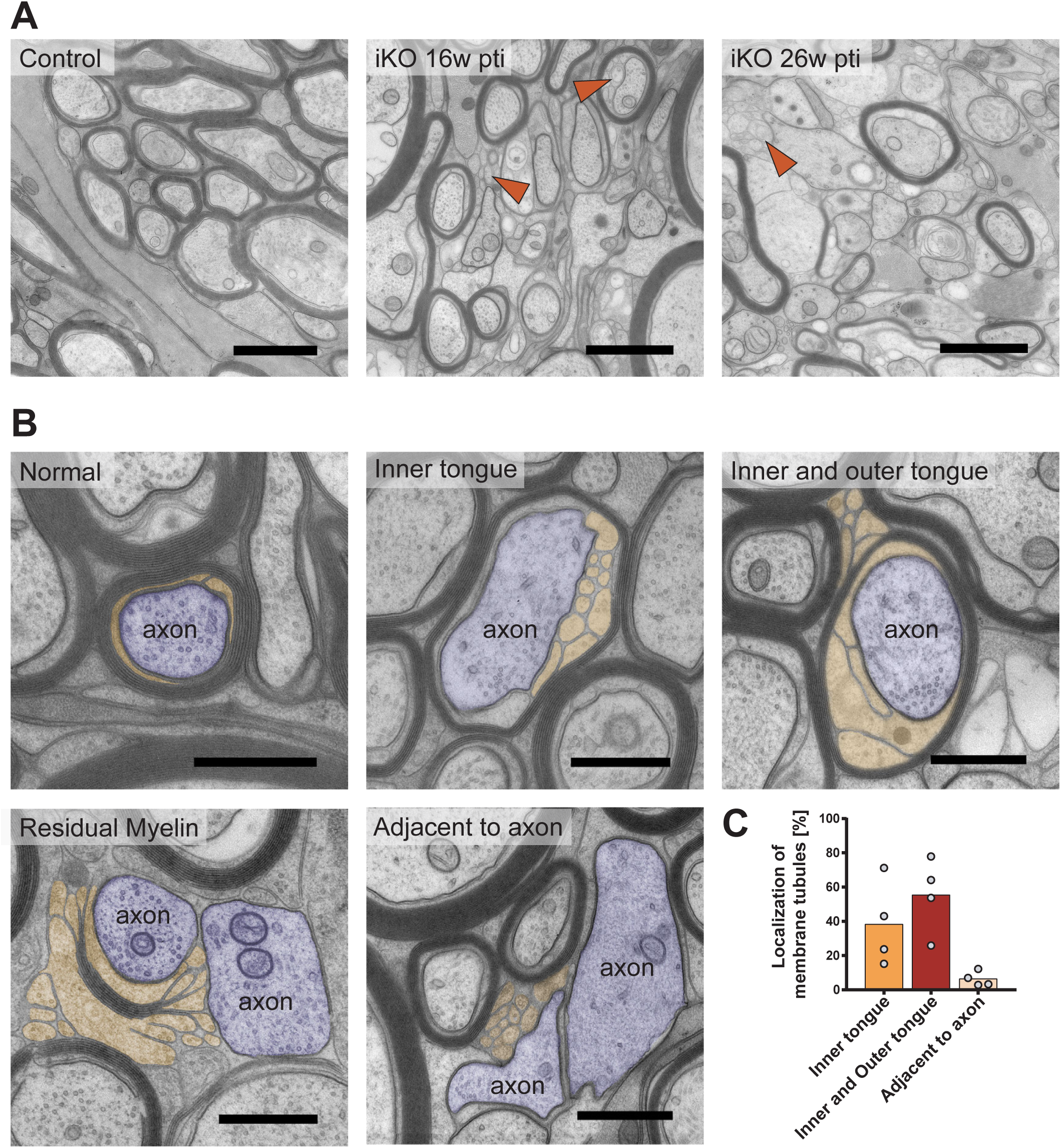
Demyelination and emergence of *shiverer*-like membrane tubules in *Mbp* iKO mice. **(A)** Electron micrographs of high-pressure frozen optic nerve showing progressive demyelination. Arrowheads indicate shiverer-like tubules. **(B)** Illustration of myelin pathology: Membrane tubules (colored in yellow) emerge at the inner tongue of iKO myelin. Tubulations at the outer tongue of a myelinated axon (colored blue) are found associated with tubulations also at the inner tongue. At places where most compact myelin is lost, membrane tubules loop out and leave a partially demyelinated axon behind. Tubules are also found next to demyelinated axons. **(C)** Quantification of the occurrence of membrane tubules. Preparation by high-pressure freezing (HPF) and freeze substitution (FS), optic nerve 16 weeks pti, n=4 iKO animals, 4 random sampled micrographs covering in total 1.600 μm^2^ were used for quantification (one-way Anova with Tukey’s multiple comparison test). Scale bars (A) 1 μm; (B) 500 nm

We quantified this phenotype on electron micrographs at the indicated time points and discovered that the loss of compact myelin apparently showed a transition phase characterized by these membrane tubules resembling *shiverer*-like myelin membranes (“pathological appearing myelinated axons” in Fig. 4A). First these membrane tubulations appear, followed by disruption and detachment of the myelin sheath from the axon, ultimately leading to almost complete demyelination. 52 weeks pti 70% of all axons were non-myelinated (Fig. 4A). By immunoelectron microscopy of optic nerve cryosections 26 weeks pti (Fig. S7), s*hiverer*-like membranes were devoid of MBP but clearly labeled for the major compact myelin protein PLP indicating that these membranes were indeed of oligodendrocytic origin.

**Figure 4:**
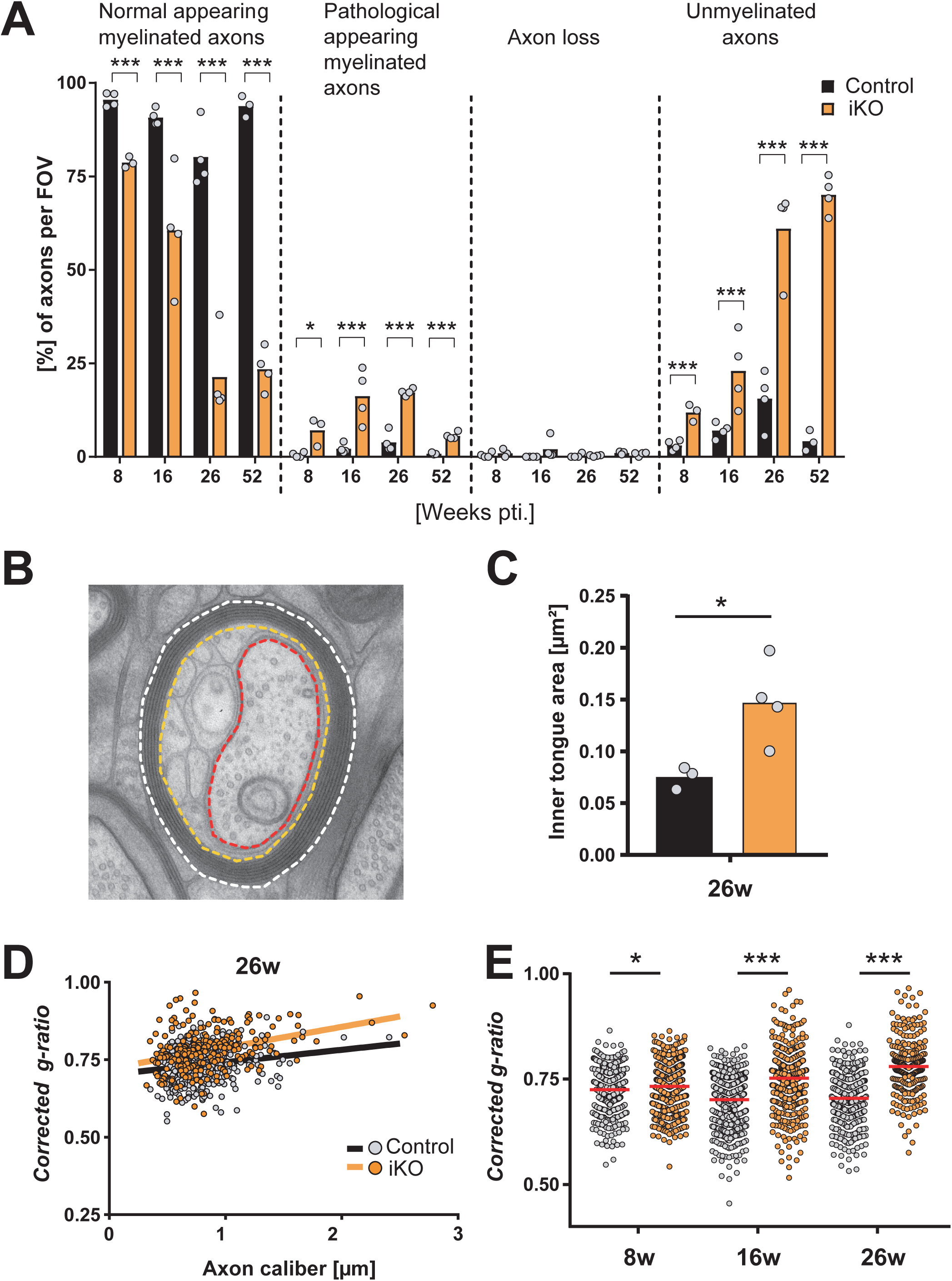
Thinning and loss of compact myelin after *Mbp* ablation. **(A)** Quantification of phenotypes at the indicated time points. Analysis was performed on optic nerve cross sections on a total area of >330 μm^2^ with >200 axons per animal, all axons in the imaged area were counted (two-tailed unpaired t-test, p<0.05 (*), p<0.01 (**), p<0.001 (***). **(B)** Illustration of corrected g-ratio measurement. Three lines are drawn for the area measurement: the outline of the fiber (stippled white line), outline of the inner border of the compact myelin (orange) and the axon (red). The area of the inner tongue was subtracted from the total fiber area before calculation of the diameter (see STAR methods). **(C)** Inner tongue area is increased in iKO compared to control in optic nerve 26 weeks pti. Measured on TEM cross sections, at least 150 axons per mouse were analyzed. T-test (p<0.05 (*) **(D)** Scatterplot depicting the corrected g-ratios at 26 weeks pti, 150 axons per mouse were analyzed. **(E)** Corrected-ratio measurements reveal an progressive decrease in compact myelin at the indicated time points, axon calibers pooled (Kolmogorow-Smirnow-Test (p<0.05 (*), p<0.01 (**), p<0.001 (***))

Interestingly, labeling density of MBP on compact myelin membrane profiles was similar between control and iKO, but the compact myelin surface area was significantly smaller in the iKO 26 weeks pti (Fig. S7C-F), in agreement with the biochemical composition of the isolated myelin fraction as shown above. Moreover, the emerging PLP-positive membrane tubules indicated ongoing myelin membrane synthesis that in the absence of MBP fails to form compact myelin.

To determine whether the loss of MBP results in thinning of the compact myelin sheath, we measured the area of the myelinated fiber and the area covered by the tubulated inner tongue and the axon (Fig. 4B). The ratio of the calculated axonal diameter divided by the calculated myelinated fiber diameter after removing the inner tongue area, which we called “corrected g-ratio”, is then plotted against the axon caliber (Fig. 4D). This corrected g-ratio is a measure for the thickness of the remaining compact myelin taking into account the enlarged tubulated inner tongue area in the iKO (Fig. 4C). At 26 weeks pti the corrected g-ratio scatter plot showed an upshift of the cloud that indicates myelin thinning (Fig. 4D). Indeed, by analysis of the myelin thickness at 8, 16 and 26 weeks pti in the optic nerve we could quantify significant myelin loss by thinning of the myelin sheath (Fig. 4E). The observed increase in the amount of non-myelinated axon profiles (Fig. 4A) indicated a loss of myelin also by shortening of internodes. To better understand the process of myelin thinning, the emergence of the *shiverer*-like membranes and internode shortening we applied 3D visualization by serial block face imaging using focused ion beam-scanning electron microscopy (FIB-SEM) (Fig. 5A) and Supplementary movie 1). The 3D reconstruction illustrates a shortened internode and a residual patch of compact myelin. By measuring the total length of myelin covering an individual axon within the imaged volume, a myelin-coverage [%] could be derived from 3D volumes at 16 weeks (n=2) and 26 weeks pti (n=1) (Fig. 5B). This measurement revealed a reduction in myelin coverage by 25% at 16 weeks pti and by 70% at 26 weeks pti. This result is in agreement with the reduced number of myelinated axons by approx. 23% (16 weeks pti) and 60% (26 weeks pti) which was determined on thin sections (Fig. 4A). To determine the time course of myelinated internode loss in our model we counted the number of myelinated axons (normally myelinated and with pathological phenotype) per area at the time points 0, 8, 16, 26, 40 and 53 weeks pti and normalized the result to control (Fig 5C). Since the amount of myelinated axons at the time point 8 weeks pti was unchanged and the reduction of myelinated axons reached the minimum at 26 weeks pti, determined a 50% loss of myelinated internodes within approx. 20 weeks pti (regression line in Fig 5C).

**Figure 5:**
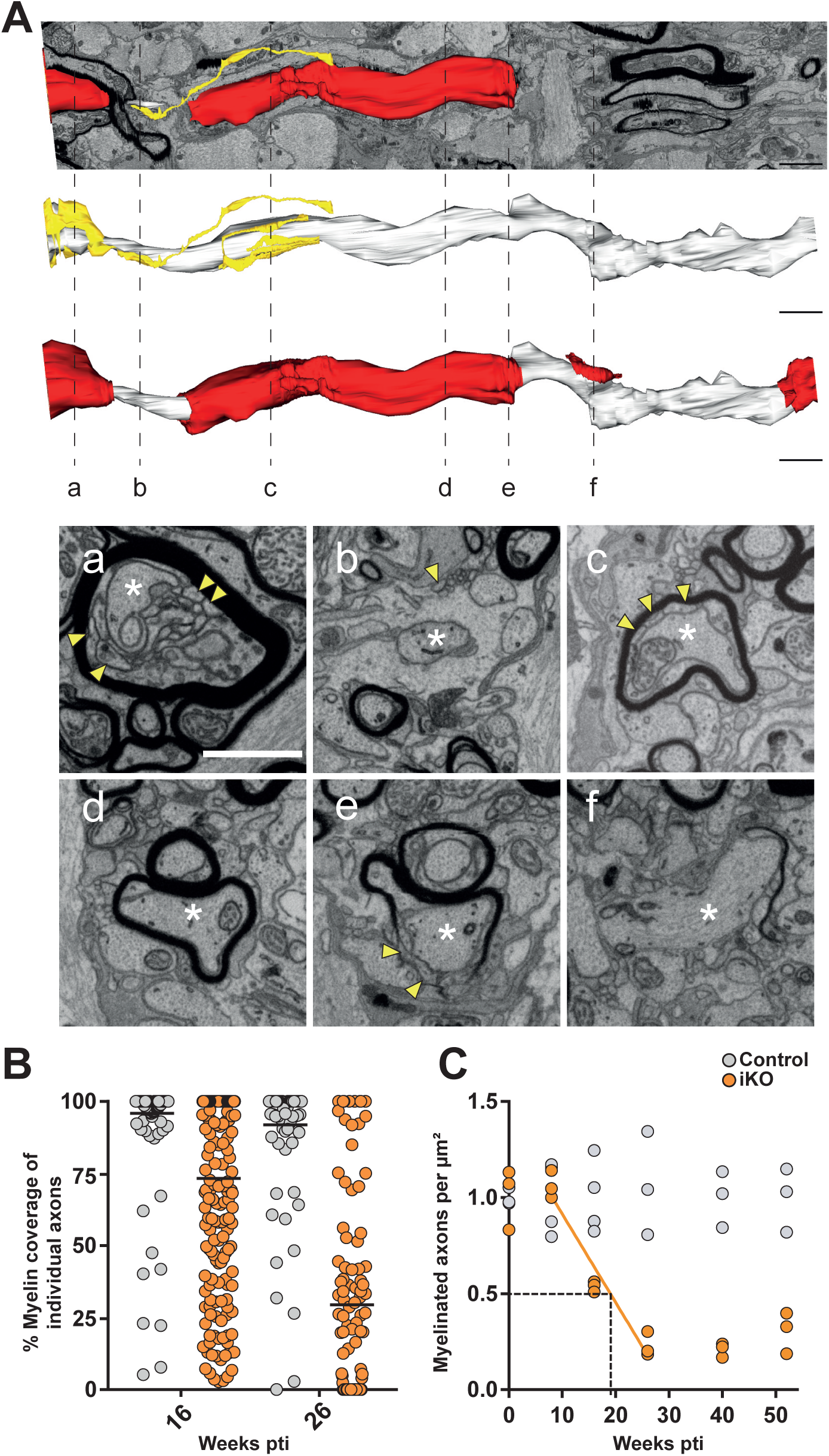
Demyelination by internode shortening in *Mbp* iKO mice. **(A)** 3D reconstruction of an image stack acquired by focused ion beam-scanning electron microscopy (FIB-SEM) at 26 weeks pti in the optic nerve of an iKO mouse (shown in Supplementary movie 1): Yellow: non-compact myelin tubules, white: axon, (white asterisk) red: myelin. At stippled lines the indicated corresponding image from the stack is shown. Yellow arrowheads point at myelin tubules. Internodes are shortened and fragmented. **(B)** Quantitation of myelin coverage on 3D volumes of (n=2) 16 weeks pti and (n=1) 26 weeks pti with >90 axons per mouse in percentage of axonal length within the FIB-SEM volume that is myelinated. **(C)** Myelinated axons counted per area on TEM images and normalized to control (n=3). The regression line indicates 50% loss of myelinated axons within 19-20 weeks pti. At 26 weeks pti demyelination is maximal. Scale bars: (A) 2 μm; (a) 1 μm

Importantly, 3D visualization revealed a number of additional aspects: The *shiverer*-like membrane processes are indeed membrane tubules and emerge at the inner tongue (Fig. 5Aa and Ac and Supplementary movie 1) and also occur at the juxtaparanode and the paranode (Fig. 6A and B) Supplementary movie 2). The remaining myelin sheaths appear fragmented and transformed into myelin tubules leaving behind some residual patches of compact myelin (Fig. 5Af). In addition to the detachment of paranodal loops, membrane tubules emerged at the paranode and often lost contact to the axon as shown in Fig. 5A as segmented structures in yellow. For comparison, we analyzed a FIB-SEM data stack from *shiverer* optic nerve at the age of 10 weeks when they reach the clinical end stage (Supplementary movie 3). Indeed the arrangement of tubular oligodendrocyte processes resembled the myelin tubules observed in the iKO. This finding supports our concept, that the induced *Mbp* knockout is gradually transforming the normal myelin sheath into shiverer myelin tubules by integration of newly synthetized material.

**Figure 6:**
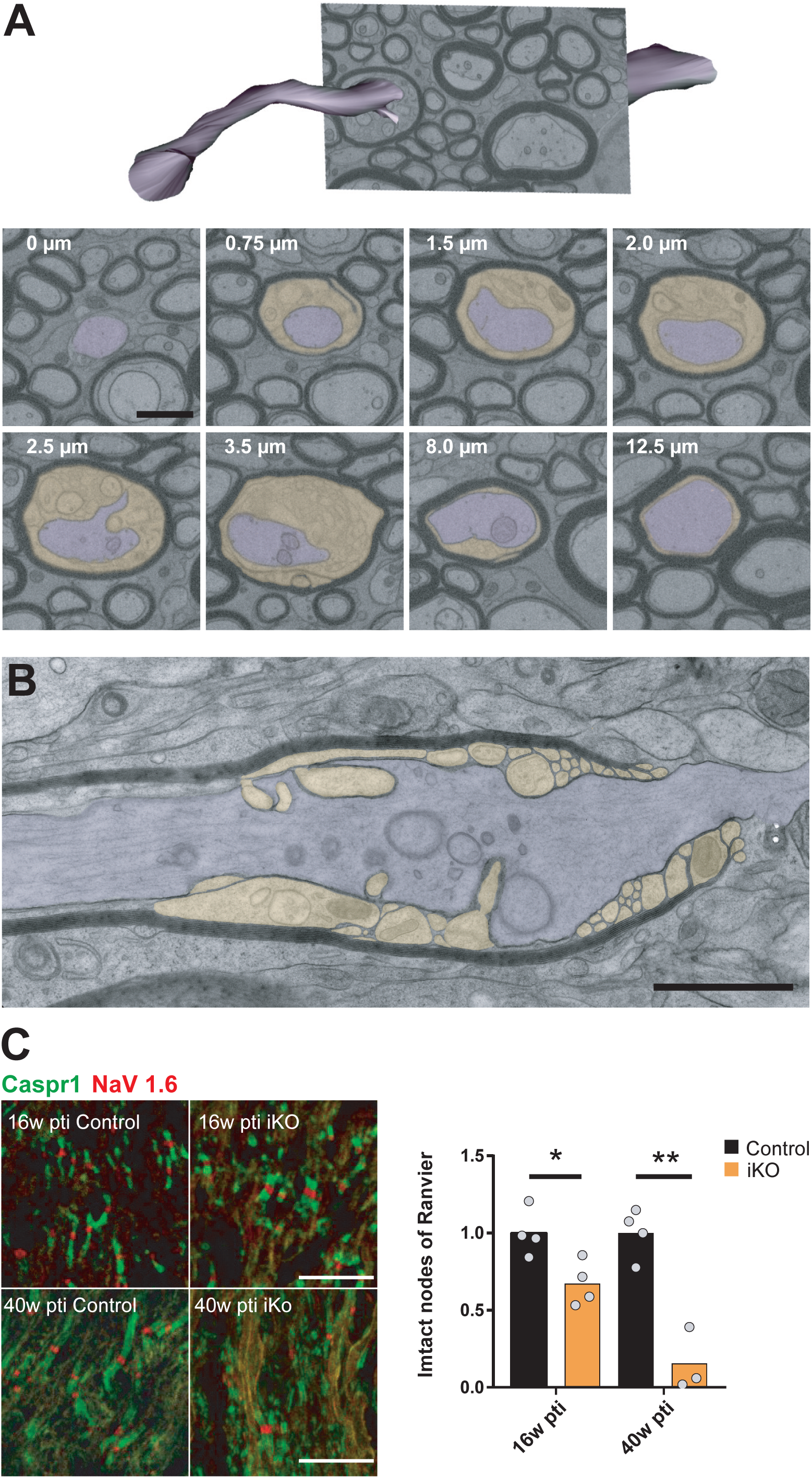
Juxtaparanodal myelin tubulation and loss of nodal organization. **(A)** Segmentation of axon and myelin tubules in an image stack acquired by FIB-SEM in optic nerve of iKO 26 weeks pti (shown in Supplementary movie 2). Distance along the internode as indicated in the images. Membrane tubules emerge at the juxtaparanode (0.75 μm-3.5 μm) while most of the internode is unaffected. **(B)** Longitudinal TEM section reveals the juxtaparanodel localization of the tubules and the detachment of the paranodal loops. **(C)** Confocal light microscopy of immunofluorescence staining of the nodal marker NaV1.6 and paranodal marker Caspr1 on optic nerve cryosections reveals loss of functional nodes of Ranvier (two-tailed unpaired t-test, p<0.05 (*), p<0.01 (**)). Scale bars: 500 nm (A, B) 10 μm (C)

To assess the nodal phenotype, we quantified nodes of Ranvier on longitudinal optic nerve cryosections after immunofluorescent staining of Caspr1 and NaV1.6 (Fig. 6C). Already at 16 weeks pti a significant loss of intact nodes was detectable which was even more pronounced at the late time point of 40 weeks pti. The loss of compact myelin in iKO mice and the accumulation of myelin tubules directly affected both the paranodal integrity and the nodal organization. Abundance changes of node proteins and others involved in axon-glia interaction detected by the proteome analysis support this evidence of disturbed node organization (Fig. 1I). We conclude that maintenance of myelin compaction by continuous MBP synthesis is essential for paranodal maintenance and node integrity.

### Myelin outfoldings, local myelin thinning and occurrence of myelinoid bodies

Maintenance of a myelin sheath probably requires a balanced input and output. We addressed the question which indications of myelin disposal are detectable and whether these are visible at specific sites. Indeed, in addition to the biosynthetic input we could also observe evidence of myelin removal in the 3D data sets obtained by FIB-SEM in optic nerve samples. Redundant myelin occurs in outfoldings in the form of large sheets of myelin extending into the vicinity and often wrapping around neighboring axons (Fig. S8A and A’). Some of the smaller outfoldings appeared as protrusions into adjacent astrocytes (Fig. S8B and B’). Microglia cells are involved in clearing of myelin debris in demyelinating conditions as well as recycling of aged myelin (Safaiyan et al., 2016, Hill et al., 2018). As expected we observed microglia containing typical lysosomes which occasionally showed close contact to myelin protrusions (Fig S8C and C’).

As shown in Fig. 4 the average myelin thickness decreased progressively after induced *Mbp* ablation. In the 3D FIB-SEM data stacks we observed that within one internode the same myelin sheath can vary remarkably in thickness (Supplementary movie 4). Moreover, we found myelinoid bodies budding at the abaxonal myelin and others protruding into the inner tongue (Fig. 7A and B and Supplementary movie 4) and also in between myelin lamellae. Myelinoid body formation at the inner tongue could indicate local myelin breakdown and uptake by the oligodendrocyte for reutilization. As we know from the NanoSIMS analysis, these structures showed a similar ^13^C to ^12^C ratio like compact myelin with little integration of newly synthetized ^13^C labeled protein. To analyze this in more detail we quantified the occurrence of myelinoid bodies in FIB-SEM data stacks over the axonal length and found a significant increase in the induced knock out at 16 weeks pti (Fig. 7C). We conclude that myelinoid body formation is enhanced after *Mbp* ablation.

**Figure 7:**
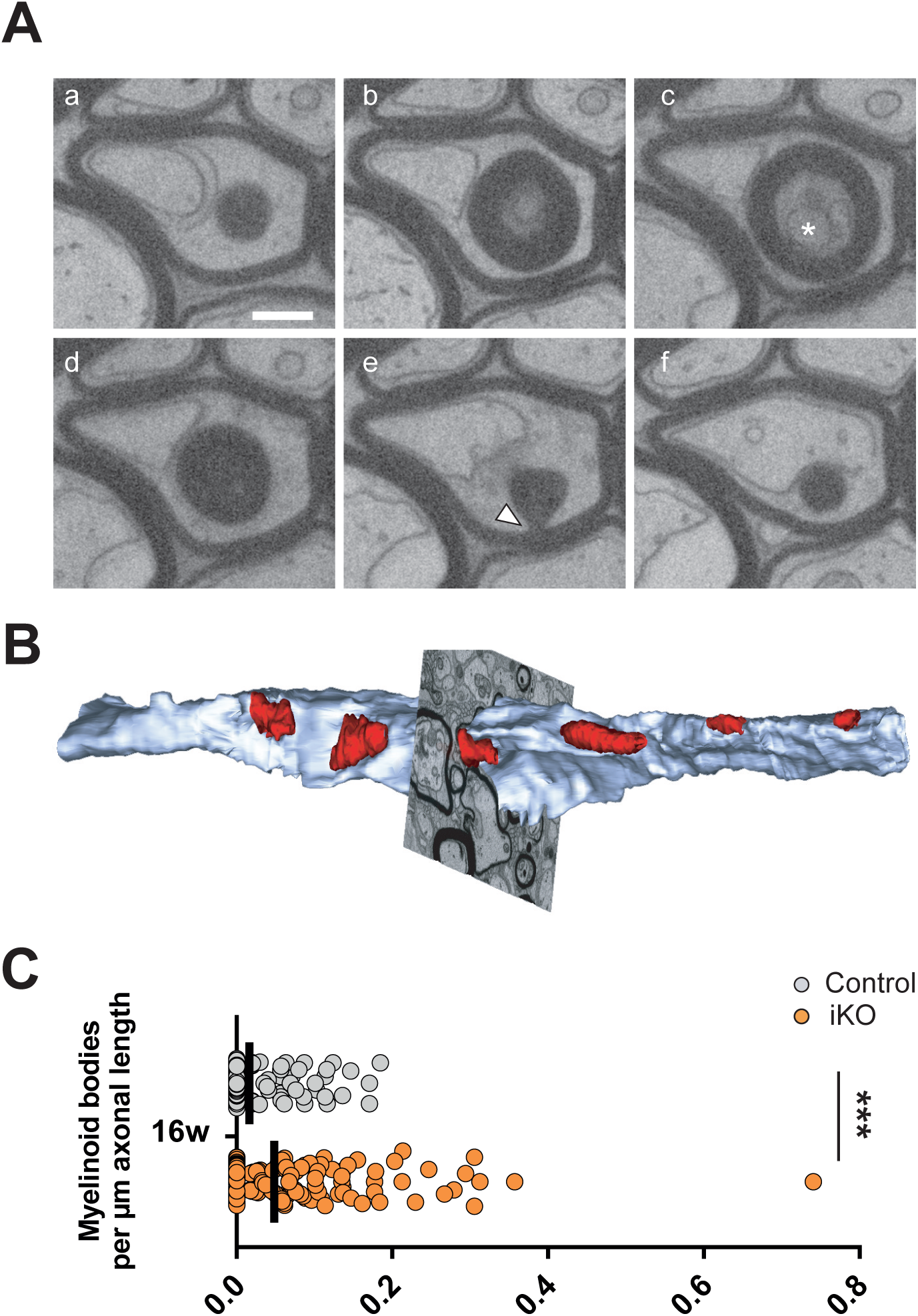
Myelinoid bodies at the inner tongue. **(A)** Electron micrographs selected from a FIB-SEM image stack reveal the presence of a myelinoid body (asterisk) at the inner tongue 16 weeks pti in high-pressure frozen optic nerve. This myelinoid body is connected to the myelin sheath (indicated by arrowhead) **(B)** Segmentation of myelinoid bodies at the inner tongue from a FIB-SEM image stack (shown in Supplementary movie 4) acquired 26 weeks pti (blue: axon, red: myelin spheres). **(C)** The number of myelinoid bodies per μm axonal length is increased 16 weeks pti on the level of individual axons. Quantification of one 3D volume with >90 axons per mouse (Kolmogorow-Smirnow-Test, wt: n=3, iKO n=2, p < .001 (***)). Scale bar: 500 nm

Based on our results we propose a mechanism of myelin turnover and renewal by which newly synthetized myelin membrane is incorporated into the sheath predominantly in the adaxonal non-compact myelin compartment at the inner tongue. Furthermore we present indications that myelin removal is not only mediated by microglia and astrocytes, but could also involve oligodendrocyte–intrinsic mechanisms by myelinoid body formation at the inner tongue.

The above described progressive loss of myelin internodes, mediated by the integration of new MBP-deficient non-compact myelin membrane and the removal of aged myelin allowed us to estimate the half-replacement time of a myelin internode in the adult optic nerve. We found that internodes shorten by 50% within 20 weeks in our mouse model. We could show that mature myelin sheaths are exceptionally stable but require a constant renewal by de novo synthesis of the components including myelin basic protein to maintain the internode and the node of Ranvier.

## Discussion

It has been recently shown that mature oligodendrocytes persist life-long in mice as well as humans (Tripathi et al., 2017, Yeung et al., 2014) and that their myelinated internodes, once made, are remarkably stable with little fluctuations in length (Hill et al., 2018). Accordingly myelin proteins are characterized by slow turnover with lifetimes in the range of weeks and months (Toyama et al., 2013, Fornasiero et al., 2018). Such biochemical studies reveal the turnover rates of the different protein or lipid components of the myelin sheath. Using inducible ablation of *Plp* in adult mice a 50% reduction of PLP within 6 months was determined (Lüders et al., 2019). One major obstacle for studies of myelin renewal is the characteristic of the myelin sheath to exclude proteins with bulky fluorescent tags (Aggarwal et al., 2011). Therefore, we designed a different strategy to make newly synthetized myelin membrane visible by exploiting the unique property of MBP to be essential for myelin compaction. We addressed two related questions: What are the morphological consequences of depleting the replenishment of myelin and what is the time-course of the emerging demyelination. Besides the effect of being able to identify new myelin by the non-compacted appearance as *shiverer*-like membranes the deletion of *mbp* in the adult mouse by targeting mature oligodendrocytes provided several major findings:

First, the reduction in MBP protein levels in the brain to 50% within 77 days (11 weeks) as calculated from our immunoblot analysis matches with the measured half-replacement time of 70 days in the cortex (Fornasiero et al., 2018) indicating normal MBP turnover under these knockout conditions. Second, MBP showed little lateral mobility, since the residual myelin fraction 26 weeks after induction contained control levels of MBP and because of the unchanged labeling density of MBP in immunoelectron microscopy on the compact myelin that remains in the iKO. Third, as already known from *shiverer* mice, oligodendrocytes differentiate without MBP expression (Bu et al., 2004). Here we show that also adult loss of MBP does not impair oligodendrocyte long-term survival. This was the key requirement for this study of internode turnover *in vivo*, since the depletion of MBP after a functional myelin sheath had been generated allowed us to detect *shiverer*-like membranes as indicators of newly synthetized myelin membrane.

We included optic nerves from 10 weeks old *shiverer* mice in the study to better understand the observed changes in the iKO on structural and proteomic level. As shown previously axons are wrapped in *shivere*r by fine tubular oligodendrocytic processes which often terminate in loops and also meander among axons without forming sheaths (Rosenbluth, 1980). We concluded that the lack of continuous *Mbp* expression in the iKO mice indeed results in a slow transformation from the previously fully myelinated nerve into a *shiverer* phenotype by replacement of the myelin membranes, since we observed similar structures in the iKO. This transformation also brought about neuropathological consequences as observed in the proteome.

We identified the inner tongue and more specifically the juxtaparanode as a metabolically active site where newly made components are integrated into the myelin sheath. The inner tongue was already determined as the growth zone in developmental myelination (Snaidero et al., 2014, Stadelmann et al., 2019). As supported by our isotope mapping experiment these cytoplasm rich compartments of the myelin sheath are the most likely places where newly synthetized components are entering the myelin sheath also in adulthood. We note that we cannot formally exclude that the outer tongue also plays an active role in myelin renewal. In our mouse model distinguishing tubulated left-overs of diminished internodes from potentially tubulated outer tongues was challenging, as the membrane tubules often lost their association with the respective axon. Since inner and outer tongue tubulations were mostly found in the same myelin sheath we conclude that the outer tongue tubules occurred closer to the end of the internode, probably representing retracted loose paranodal loops.

The internodal shortening by the integration of non-compact *shiverer*-membranes at the paranode and juxtaparanode resulted in membrane tubulation and an impairment of the nodal organization. More specifically, in our proteome analysis we found a concomitant reduction of the protein levels of the juxtaparanodel voltage gated potassium channel α subunit K_v_1.2 and the associated disintegrin and metalloprotease ADAM22. This enzyme is an axonal component of the juxtaparanodal macromolecular complex composed of K_v_1.2 (Ogawa et al., 2010) and cell adhesion proteins Caspr2 and contactin-2 (TAG-1) (Poliak et al., 2003, Traka et al., 2003, Horresh et al., 2008). In accordance we found a reduction of contactin-2 abundance at the late time point (40 weeks iKO) and in *shiverer*. Alterations of K_v_1.1 and K_v_1.2 channel subunit distribution were also described in *shiverer* mice (Sinha et al., 2006). However, in this study an increased abundance of K_v_1.2 was found in *shiverer* spinal cord whereas we found a decrease in the whole optic nerve proteome. Alterations of paranodal axo-glial junctions, the organization of the juxtaparanode and retraction of paranodes have been reported in the EAE model of demyelination and in MS patients (Fu et al., 2011, Howell et al., 2006, Coman et al., 2006). Different from those models of inflammatory demyelination in our model retraction of paranodes and demyelination occurred as a consequence of genetic ablation of MBP and developed very slowly without causing oligodendrocyte loss. Paranodal abnormalities without oligodendrocyte loss also occur in mice with genetic ablation of myelin galactosphingolipid synthesis (Dupree et al., 1998, Bosio et al., 1998). However, these mice suffer from dysmyelination, develop a progressive neuropathology and die prematurely. In our model, the reduction of the abundance the UDP-galactose:ceramide galactosyl-transferas (UGT8) and of several enzymes involved in isoprenoid and cholesterol synthesis determined by proteome analysis is consistent with the observed nodal phenotype. Node of Ranvier maintenance depends not only on the complete set of axo-glial interacting proteins, but also requires intact membrane microdomains composed of galactospingolipids, gangliosides and cholesterol (Poliak and Peles, 2003, Dupree et al., 1999, McGonigal et al., 2019, Susuki et al., 2007). Since MBP interacts with negatively charged lipids such as PI(4,5)P_2_, influences lipid ordering and associates with galactosylceramid and cholesterol-rich lipid rafts in mature myelin (DeBruin et al., 2005, Fitzner et al., 2006, Musse et al., 2008, Debruin and Harauz, 2007, Ozgen et al., 2014), decrease of MBP levels together with changes in its posttranslational modifications could affect membrane composition and function (Boggs, 2006). In addition, breakdown of diffusion barriers by the loss of myelin compaction in our *Mbp* iKO model could also impact paranodal and juxtaparanodal integrity. Striking morphological similarities exist between our model and the transcription factor *Nkx6-2* null mouse (Southwood et al., 2004) in the form of myelin tubules (‘vermicular-like processes’) at the inner tongue and compact myelin ‘flaps’ or stacks flanked by detached paranodal loops. However, these similar phenotypes seem to arise by different mechanisms. In case of the *Nkx6-2* null mouse dysregulated expression of paranodal proteins and defects in cytoskeletal remodeling might play important roles, whereas we explain the phenotype in the *Mbp* iKO by the continuous integration of newly formed myelin membrane and the lack of myelin compaction.

We have shown here that induced *Mbp* deletion resulted in a progressive loss of compact myelin. So, how is this compartment eliminated? In accordance with the literature of the time Hildebrand and colleagues proposed a concept of a metabolically active myelin sheath with lifelong turnover utilizing a ‘quantal’ detachment of Marchi-positive myelinoid bodies as disposal route (Hildebrand et al., 1993). These lamellated myelinoid bodies were found preferentially at the paranodes of large myelinated fibers in the cat spinal cord and inside astrocytes and microglia (Hildebrand, 1971, Persson and Berthold, 1991). When we mapped the occurrence of myelinoid bodies along axons in the optic nerve in our 3D data stacks, we could not detect a preferential localization. As already discussed in the above mentioned review (Hildebrand et al., 1993), there might be qualitative differences in myelin turnover regarding myelinoid body formation between large myelinated fibers as in the spinal cord and small caliber fibers like in the optic nerve.

As published recently, phagocytosis of myelin debris by microglia increases with age and leads to lipofuscin accumulation in the microglial lysosomal compartment (Safaiyan et al., 2016). The same study found similar effects in *shiverer* mice and also described occasional myelin uptake by astrocytes. Age-related accumulation of myelin debris within microglia was also demonstrated by *in vivo* imaging in the cerebral cortex of the mouse (Hill et al., 2018). Myelin phagocytosis by astrocytes seem to play a role in myelin remodeling under certain non-pathological conditions such as internode shortening in the frog optic nerve during metamorphosis (Mills et al., 2015). Myelin debris uptake by astrocytes was also described in different forms of white matter injury as an early event in lesion formation leading to enhanced inflammation (Ponath et al., 2017). Under conditions of myelin degeneration and oligodendrocyte damage the formation of myelinosomes was identified as early event in demyelination and lesion development (Romanelli et al., 2016). These myelinosomes show striking similarity in morphology to the myelinoid bodies we found in our model. Yet, different to the study of Romanelli and colleagues the *Mbp* iKO mouse is unlike any model of acute demyelination because of the absence of oligodendrocyte damage. In our 3D data sets we also found an increased number of myelinoid bodies inside the inner tongue resembling a ‘myelin inclusion’ described in (Romanelli et al., 2016). Such myelinoid bodies were also visible at the inner tongue under control conditions. However, this seems to be an infrequent event that easily escapes electron microscopic observation on thin sections.

A recent study demonstrated that lipid metabolism is essential for myelin integrity (Zhou et al., 2020). Here, Qki-5, a transcriptional coactivator of the PPARβ-RXR*α* complex which regulates fatty acid metabolism, was depleted by induced knock out in the adult mouse. Strikingly, rapid and severe demyelination started as early as one week after tamoxifen induction without impairing oligodendrocyte survival. In this model, the loss of myelin lipids was accompanied by conformational changes of MBP promoting dissociation from membranes and loss of myelin compaction. This study indicates that myelin lipids turn over much faster than myelin proteins.

By the application of advanced imaging methods in combination with a novel mouse model to deplete compact myelin maintenance, we visualize the myelin subcompartment at which newly synthetized myelin membranes are added. We identified the paranodal and juxtaparanodal region at the inner tongue as important site for myelin renewal in adults and found indications of myelin disposal in form of myelinoid bodies occurring abaxonally and in the inner tongue.

## Supporting information

Supplementary Figures_Meschkat et al

## Acknowledgement

We thank R. Jung, A. Fahrenholz, D. Hesse and U. Kutzke for technical support and U. Suter for the inducible *Plp*-Cre^ERT2^ mice. This work was supported by the Deutsche Forschungsgemeinschaft (DFG) (FOR2848, project 08 to W.M.) and DFG grants (WE 2720/2-2 and WE 2720/4-1 to H.B.W) and by the Cluster of Excellence and Deutsche Forschungsgemeinschaft (DFG) Research Center Nanoscale Microscopy and Molecular Physiology of the Brain (W.M., A.M.S., H.E. and K.A.N) and the ERC (advanced grant to K.A.N.), by the SFB1286/B1 to P. Agüi-Gonzalez and the SFB1286/B1 and VR (Swedish Research Council) to N. T. N. Phan.

## Author contributions

Conceptualization, W.M. and M.M.; Methodology, M.M., A.M.S., M.-T.W., K.K., O.J., L.P. P.A.-G., N.T.N.P., T.R., B.S. and S.R.; Software, S.R.; Investigation, M.M., O.J. and W.M.; Formal analysis, M.M., L.P., O.J. and P.A.-G.; Writing-Original draft, W.M.; Writing-Review and Editing Final Draft, M.M., A.M.S., O.J., S.R., H.B.W., H.E. and K.A.N.; Visualization, M.M., A.M.S. and L.P.; Supervision, W.M.; Funding Acquisition, W.M., P.A.-G., N.T.N.P., H.E. and K.A.N.

## STAR Methods

### LEAD CONTACT AND MATERIALS AVAILABILITY

Further information and requests for resources and reagents should be directed to and will be fulfilled by the Lead Contact, Wiebke Möbius. Mouse lines generated in this study are available after an MTA is signed.

### EXPERIMENTAL MODEL AND SUBJECT DETAILS

All animal experiments were performed in accordance with the German and European animal welfare laws and approved by the Lower Saxony State Office for Consumer Protection and Food Safety (license 33.19-42502-04-16/2119). All mice were housed in standard plastic cages with 1-5 littermates in a 12 h/12 h light/dark cycle (5:30 am/5:30 pm) in a temperature-controlled room (∼21°C), with *ad libitum* access to food and water. All mice used were bred under the C57BL6/N background. Experiments were carried out mostly in male and sometimes in female mice (indicated in the data). For the generation of the *Mbp*^fl/fl^ mouse line embryonic stem (ES) cells harboring a modified allele of the *Mbp* gene (*Mbp*^tm1a^) carrying a LacZ-neomycin cassette upstream of exon 1 of the classical *Mbp* locus were acquired from the European Conditional Mouse Mutagenesis Program (Eucomm). ES cells were microinjected into blastocysts derived from FVB mice and embryos were transferred to pseudo pregnant foster mothers. For the ES clone B02 germline transmission was achieved by breeding with C57BL/6N female mice. The resulting offspring harbored the Mbp-lacZ neo allele (*Mbp*^neo/neo^). The construct including the lacZ gene and a neomycin resistance cassette was excised by crossbreeding with mice expressing a FLIP recombinase (129S4/SvJaeSor*-*Gt(ROSA)26Sortm1(FLP1)Dym/J; backcrossed to C57BL6/N). Mice expressing tamoxifen inducible Cre^Ert2^ under the control of the *Plp* promotor (MGI:2663093) (Leone et al., 2003) were obtained from Ueli Suter, ETH Zurich Institute for Molecular Health Sciences, Switzerland. Control mice (Mbp^fl/fl^*Plp^*CreERT2wt*^) and inducible knockout mice (iKO) (Mbp^fl/fl^*Plp^*CreERT2+*^) were generated by breeding Mbp^fl/fl^*Plp^*CreERT2wt*^ males with Mbp^fl/fl^*Plp^*CreERT2+*^ females. For experiments control and iKO mice were used in groups of littermates of the same sex. Mice were sacrificed at the indicated time points after tamoxifen induction. Genotyping of the *Mbp* flox allele was performed by genomic polymerase chain reaction (PCR) using the following primers: *Mbp* wt fwd: 5’-GGGTGATAGACTGGAAGGGTTG *Mbp* wt rev: 3’ of LoxP site: 5’-GCTAACCTGGATTGAGCTTGC Lar3 rev: 5’-CAACGGGTTCTTCTGTTAGTCC Genotyping of the Cre^Ert2^ allele was performed using the following primers: 5’-CAGGGTGTTATAAGCAATCCC 5’-CCTGGA AAATGCTTCTGTCCG, including a primer pair for CNP as positive control: 5’-GCCTTCAAAC-TGTCCATCTC 5’-CCCAGCCCTTTTATTACCAC

## METHOD DETAILS

### Tamoxifen induction

For knock-out induction 8-9 weeks old mice were injected intraperitoneally with 1 mg tamoxifen (100 μl of 10 mg/ml tamoxifen (Sigma-Aldrich, St. Louis, MO) in corn oil (Sigma-Aldrich)) for 5 consecutive days, followed by a two-day break and 5 more days of injection as described (Leone et al., 2003). To prepare the tamoxifen solution, in a 2 ml tube, 500 μl ethanol and 500 μl corn oil were added to 50 mg tamoxifen and mixed in a tissue lyser (Qiagen, Hilden, Germany) for 10 min at 50 Hertz. After this the resulting emulsion was added to 4 ml corn oil and mixed until the solution turned clear. The solution was stored in the fridge and used within 5 days. The day after the last tamoxifen injection was considered 0 days post tamoxifen.

### Expression analyses

For the characterization of myelin gene expression, RNA from total spinal cord and brain lysates as well as corpus callosum was isolated using QIAzol (Qiagen) and the RNeasy protocols (Qiagen). Concentration and purity of RNA was evaluated using a NanoDrop spectrophotometer (Thermo Fisher Scientific, Waltham, MA, USA). Complementary DNA (cDNA) was synthesized using the Superscript III (Invitrogen, Carlsbad, CA, USA) according to the manufacturer’s protocol. Quantitative RT-PCR was performed in triplicates with the GoTaq qPCR Master Mix (Promega) on a LightCycler 480 II PCR system (Roche, Basel, Switzerland). Expression was normalized to the mean of two housekeeping genes Rps13 (Ribosomal Protein S13) and PPIA (Cyclophilin A). Relative changes in gene expression were analyzed using the 2ΔΔC(T) method (Livak and Schmittgen, 2001). Primers were designed using the Universal Probe Library form Roche Applied systems (https://www.roche-applied-science.com) and validated using NIH PrimerBlast (www.ncbi.nlm.nih.gov/tools/primer-blast/).

Expression of the following genes was *Car2* (carbonic anhydrase 2), *Cnp* (2’,3’-cyclic nucleotide 3’ phosphodiesterase), *Golli* (Golli-MBP), *Mag* (myelin associated glycoprotein), *Mbp* (myelin basic protein), *Mog* (myelin oligodendrocyte glycoprotein), *Olig2* (oligodendrocyte lineage transcription factor 2), *Pdgfa* (platelet derived growth factor alpha), *Plp* (proteolipid protein). All primer sequences are listed in Table List of primer sequences. All primers used for expression analysis were intron-spanning (5’-3’; forward - reverse).

### Table: List of primer sequences

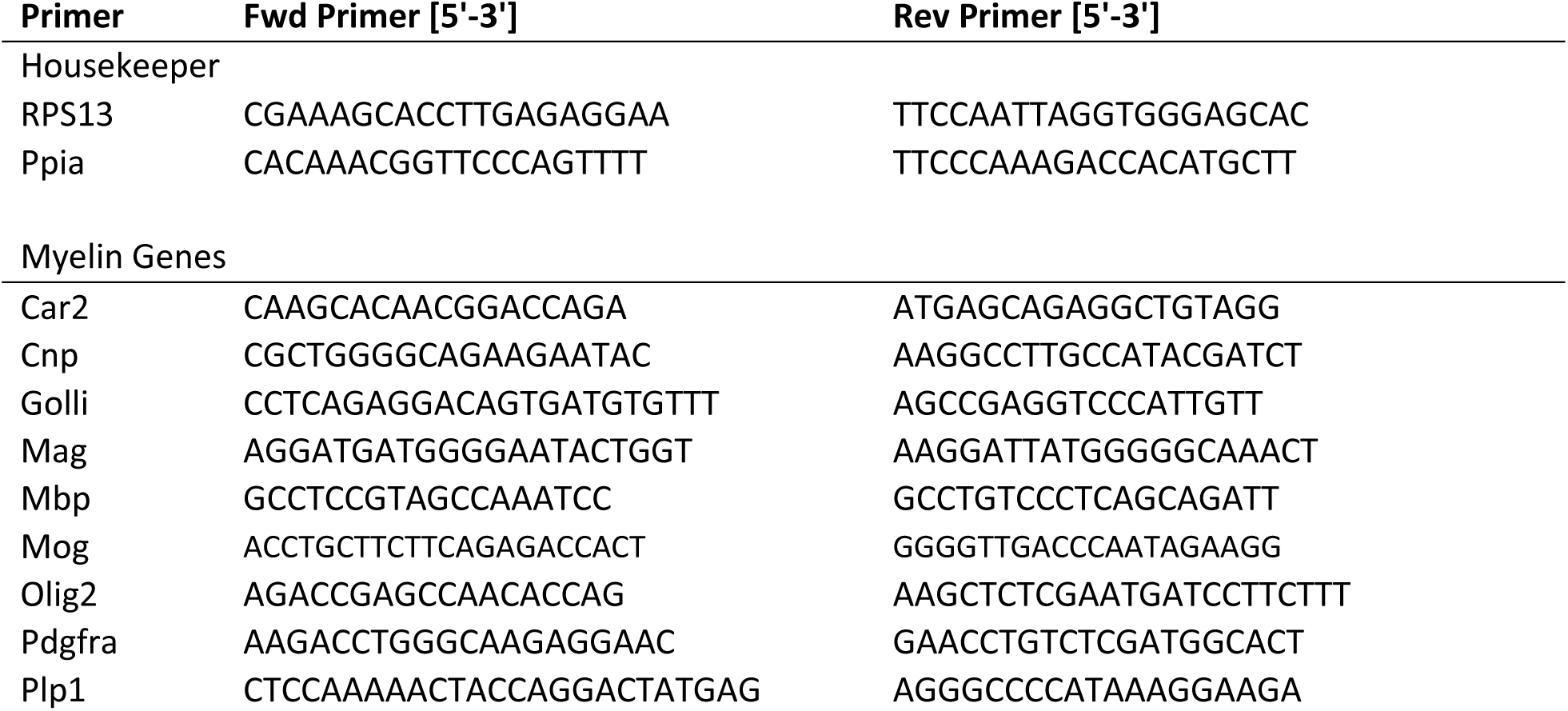

### Tissue lysis

For whole brain lysate, one hemisphere was homogenized in 4 ml modified RIPA buffer using an T-10 basic Ultra-Turrax (IKA, IKA®-Werke GmbH & CO. KG, Staufen Germany) for 20-30 sec on speed setting 3-4. The homogenate was then centrifuged at 13.000 rpm for 10 minutes at 4°C and the supernatant was transferred to a new tube. Protein concentration was determined in triplicates using the BCA protein assay kit (Pierce) according to the manufacturer’s manual.

### Myelin purification and immunoblotting

Purification of a light*-*weight membrane fraction enriched for myelin was performed as previously described (Erwig et al., 2019a). For myelin purification, half brains of three male control and iKO mice each at 26 weeks post tamoxifen were homogenized in 0.32 M sucrose. Purified myelin was taken up in 1⨯ TBS with protease inhibitor (Complete Mini, Roche). For lysate analysis half brains of three Ctrl and iKO mice each at 8, 16 and 26 weeks post tamoxifen were homogenized in modified RIPA buffer (1x TBS, 1 mM EDTA, 0.5% [w/v] Sodium deoxycholate, 1.0% [v/v] Triton X-100, cOmplete™ Mini protease inhibitor (Roche) using a T-10 basic Ultra-Turrax. Protein concentration was measured using the DC protein assay (BioRad Laboratories, Hercules, CA) according to the manufacturer’s guidelines.

Immunoblotting was performed as previously described (Kusch et al., 2017). Purified myelin (0.5 μg for PLP/DM20; 2.5 μg for MBP) was separated on SDS*-*polyacrylamide gels (15% for PLP/DM20 and MBP) and blotted onto PVDF membranes (Hybond; Amersham) using the XCell II Blot Module (Invitrogen). Primary antibodies were incubated overnight at 4 °C in 5% milk in TBS with 0.1% Tween 20. HRP coupled secondary antibodies α*-*rabbit*-*HRP (Dako) or α*-*mouse*-*HRP (Dako) (1:10000) were incubated in 5% milk in TBS with 0.1% Tween 20 for 1 hr at RT and detected using a CHEMOSTAR ECL & Fluorescence Imager (Intas). Quantification was performed in ImageJ (Fiji) (Schindelin et al., 2012) using actin or fast-green total protein as loading control, graphs were plotted using GraphPad Prism 7.0. Statistical evaluation was performed using a two-tailed unpaired t-test (GraphPad Prism 7.0) per individual time point (iKO vs age-matched control). Levels of significance were displayed as p < .05 (*), p < .01 (**), and p < .001 (***).

### Proteome analysis of whole optic nerve lysates

To investigate the systemic response to the loss of MBP and compact myelin in an adult animal we performed proteome analysis of whole optic nerves at 8W, 16W and 40W post tamoxifen. To compare the progressive loss of MBP during adulthood with the complete absence of MBP we also included optic nerves of 10 week old *shiverer* animal, a naturally occurring MBP knockout. Whole optic nerves were homogenized in 250 μl ice cold modified RIPA buffer (1x TBS, 1 mM EDTA, 0.5%[w/v] sodium deoxycholat, 1.0%[v/v] Triton X-100, c0mplete^*™*^ Mini protease inhibitor (Roche Diagnostics)) using Teflon beads and a Precellys 24 tissue homogenizer (Bertin instruments, France). Homogenization was carried out at a speed of 5500 rpm for 2×10 seconds. Samples were then centrifuged at 13.000 rpm at 4°C and the supernatant was collected to reduce undissolved tissue and cellular nuclei. The protein concentration of the supernatant was determined in triplicates using the BCA protein assay kit (Pierce). For quality control 0.5 μg protein of each sample were loaded on a gel and SDS-PAGE with subsequent silver staining of the gel was performed to visualize protein bands.

Supernatant fractions corresponding to 10 μg protein were subjected to in-solution digestion by filter-aided sample preparation (FASP) according to a protocol modified for processing of purified myelin as recently described in detail (Erwig et al., 2019a, Siems et al., 2020). Optic nerve protein samples were lysed and reduced in lysis buffer (7 M urea, 2 M thiourea, 10 mM DTT, 0.1 M Tris pH 8.5) containing 1% ASB-14, followed by dilution with ∼10 volumes lysis buffer containing 2% CHAPS to reduce the ASB-14 concentration. Samples were loaded on centrifugal filter units (30 kDa MWCO, Merck Millipore), detergents were removed with wash buffer (8 M urea, 10 mM DTT, 0.1 M Tris pH 8.5), proteins were alkylated with 50 mM iodoacetamide in 8 M urea, 0.1 M Tris pH 8.5, and excess reagent was removed with wash buffer. After buffer exchange with 50 mM ammonium bicarbonate (ABC) containing 10 % acetonitrile, proteins were digested overnight at 37°C with 400 ng trypsin in 40 μl of the same buffer. Recovered tryptic peptides were spiked with 10 fmol/μl of yeast enolase-1 tryptic digest standard (Waters Corporation) for quantification purposes and directly subjected to LC-MS analysis using nanoscale reversed-phase UPLC separation (nanoAcquity system, Waters Corporation) coupled to quadrupole time-of-flight mass spectrometry with ion mobility option (Synapt G2-S, Waters Corporation). Peptides were trapped for 4 min at a flow rate of 8 μl/min 0.1% TFA on Symmetry C18 5 μm, 180 μm × 20 mm trap column and then separated at 45°C on a HSS T3 C18 1.8 μm, 75 μm × 250 mm analytical column over 140 min at a flow rate of 300 nl/min with a gradient comprising two linear steps of 3-40% mobile phase B in 120 min and 40-60% mobile phase B in 20 min (A, water/0.1% formic acid; B, acetonitrile/0.1% formic acid). Mass spectrometric analysis was performed in the ion mobility-enhanced data-independent acquisition mode with drift time-specific collision energies (referred to as UDMSE) as introduced by Distler and colleagues (Distler et al., 2014, Distler et al., 2016) and adapted by us to synaptic protein fractions (Ambrozkiewicz et al., 2018) and purified myelin (Siems et al., 2020). For the correct quantification of highest abundance proteins such as PLP and MBP, all samples were re-run in a data acquisition mode without ion mobility separation of peptides (referred to as MS^E^) to provide a maximal dynamic range at the cost of proteome coverage (Siems et al., 2020). Continuum LC-MS data were processed for signal detection, peak picking, and isotope and charge state deconvolution using Waters ProteinLynx Global Server (PLGS) version 3.0.3. For protein identification, a custom database was compiled by adding the sequence information for yeast enolase 1 and porcine trypsin to the UniProtKB/Swiss-Prot mouse proteome (release 2018-11, 17001 entries) and by appending the reversed sequence of each entry to enable the determination of false discovery rate (FDR). Precursor and fragment ion mass tolerances were automatically determined by PLGS and were typically below 5 ppm for precursor ions and below 10 ppm (root mean square) for fragment ions. Carbamidomethylation of cysteine was specified as fixed and oxidation of methionine as variable modification. One missed trypsin cleavage was allowed. Minimal ion matching requirements were two fragments per peptide, five fragments per protein, and one peptide per protein. The FDR for protein identification was set to 1% threshold.

Per condition (8W pti, 16W pti, 40w pti, shiverer), optic nerve fractions from four animals per genotype (Ctrl, iKO) were processed with replicate digestion, resulting in two technical replicates per biological replicate and thus in a total of 16 LC-MS runs to be compared.

### Antibodies

Antibodies against the following antigens were used: APP (Chemicon MAB348), CAII (gift from Said Ghandour (Ghandour et al., 1979)); CD3 (Abcam ab11089), GFAP (Novocastra NCL-GFAP-GA5), Iba1 (Abcam ab5076), MAC3 (Pharmigen 553322); MBP (this study: for generation of MBP antisera, rabbits were immunized with the intracellular peptide 105-115 of the 21.5kDa isoform of mouse MBP (CQDENPVVHFFK). Anti-MBP antibodies were purified by affinity chromatography. The epitope is conserved in human and rat.), Olig2 (gift from Charles Stiles/John Alberta, DF308, (Sun et al., 2003)), Caspr1 (Neuromab K65/35), Nav 1.6 (Alomone ASC-009), PDGFRA (Cell Signaling 3174), PLP1 (polyclonal rabbit, A431 (Jung et al., 1996)), MAG (Millipore Ab1567), MOG (gift from Christopher Linnington (Linnington et al., 1984)), Actin (Millipore Mab 1501)

### Immunohistochemistry

For Caspr1, Nav 1.6 and MBP labeling, mice were anaesthetized with Avertin (Weil et al., 2019) and flushed with Hanks balanced salt solution (HBSS) followed by perfusion with 4% PFA in 0.1M phosphate buffer. Brain and optic nerve were dissected. Optic nerves were postfixed in 4% PFA for 10 minutes and prepared for cryosectioning. Brain was postfixed in 4% PFA for 24 hours and prepared for paraffin embedding. Slide-mounted optic nerve cryo sections (9 μm) were air-dried at RT and washed three times 10 min in PBS followed by permeabilization for one hour in 0.4% Triton-100 in PBS. Sections were blocked for 1 h in 1% fetal calf serum, 1% bovine serum albumin and 1% fish skin gelatin in 1x PBS. The primary antibodies were then diluted in the blocking solution and incubated overnight at 4°C. On the next day, slides were washed thrice with 1x PBS for 15 min each and incubated with the fluorescently labeled secondary antibody for 1 hour at RT. After the incubation the slides washed thrice with 1x PBS for 15 min each, incubated with DAPI for 20 min and washed again before being mounted using Aqua-Poly/Mount (Polysciences). Sections were stored at 4°C until imaging. Brain hemispheres were embedded in paraffin, sectioned coronally and labeled chromogenic for CD3, GFAP, APP, MAC3 and with fluorescent antibodies for CAII, Olig2 and PDGFRa as essentially described in (Stumpf et al., 2019a, Stumpf et al., 2019b). TUNEL staining of paraffin fimbria sections for quantification of apoptotic cells was performed according to the manufacturer’s protocols (DeadEnd™ Colorimetric TUNEL System, Promega). Imaging of 2 fimbria (GFAP, MAC3, APP, Olig2, CAII, PDGFRA) or 6 fimbria (CD3) per animal was performed on a bright*-*field light microscope (Zeiss AxioImager Z1 with Zeiss AxioCam MRc camera) with the following magnifications: 20x (CD3, GFAP, MAC3, Olig2, CAII, PDGFRA), 40x (APP), and 100x (representative images for display in the figures). Image analysis and quantification of markers was performed in Fiji (Schindelin et al., 2012) using a custom made thresholding macro (GFAP, MAC3) or counted manually using the cell counter plugin (CD3, APP) in Fiji.

### EDU labeling

Mice at the time point 40 weeks after tamoxifen administration received 5-ethynyl-2′-deoxyuridine (EdU) in drinking water 0.2 mg/ml for 3 consecutive weeks. Mice were killed for analysis 3 weeks after the final EDU administration. Labeling of EDU positive cells on paraffin sections of fimbria was performed using the Click-iT™ Plus EdU Cell Proliferation Kit (Thermo Fischer Scientific) according to the manufacturer’s protocol followed by fluorescent co-labeling for CAII, Olig2 or PDGFRA as described.

### Confocal microscopy

Confocal images were acquired on a SP5 confocal microscope (Leica Microsystems). Fluorescent signals were imaged sequentially to avoid cross bleeding using an HCX PL APO lambda blue 63.0×1.20 WATER UV objective. The following laser lines were used Argon Laser at 488 nm and 514 nm was used to excite Alexa 488 and Alexa 555 respectively. A HeliumNeon (HeNe) laser at 633 nm was used to excite Alexa 633 and Alexa 647. The confocal software LasAF was used for image acquisition. Images were saved as .lif and quantified using Fiji. The number of nodes of Ranvier was quantified in 9 μm cryostat sections of optic nerves 16W and 40W pti in at least 3 FOV with 240×240 μm length.

### Electron microscopy

Sample preparation of optic nerves by high-pressure freezing (HPF) and freeze substitution (FS) for transmission electron microscopy was performed as described (Weil et al., 2019). Mice were killed by cervical dislocation and optic nerves were dissected and placed into an HPF specimen carrier with an indentation of 0.2 mm. The remaining volume was filled with 20% polyvinylpyrrolidone (Sigma*-*Aldrich, P2307*-*100G) in PBS. Samples were cryo immobilized and fixed using a HPM100 (Leica) and freeze substituted using a Leica AFS (Leica Microsystems, Vienna, Austria) and embedded in Epon resin according to the protocol for optic nerves (Weil et al., 2019). Tissues for conventional fixation used for quantification of corrected g-ratios and phenotype counting were dissected and immersion fixed for at least 24 h in 4% paraformaldehyde (PFA) and 2.5% glutaraldehyde in 0.1 M PB (containing 109.5 mM NaH_2_PO_4_·H_2_O, 93.75 mM Na_2_HPO_4_·2H_2_O, and 86.2 mM NaCl), contrasted with osmium tetroxide, dehydrated and Epon embedded as described (Weil et al., 2019). Ultrathin sections (50 nm) were cut using a UC-7 ultramicrotome (Leica Microsystems, Vienna, Austria) and contrasted with UranyLess™ (Science Services, Munich, Germany) for 30 min. Samples were imaged using a LEO 912AB Omega transmission electron microscope (Carl Zeiss, Oberkochen, Germany) with an on*-*axis 2048 x 2048*-*CCD*-*camera (TRS, Moorenweis, Germany).

Normal appearing myelinated axons, axons with visible membrane tubulations, axonal degeneration and unmyelinated axons were quantified on TEM micrographs. Quantification was performed on 3-4 animals per time point with at least 5 FOV with a total area of at least 330 μm^2^ and at least 200 axons per animal. Statistical analysis was performed between iKO and the respective control using a two-tailed unpaired t-test.

For quantification of inner tongue area and measurement of corrected g-ratio, images were analyzed using Fiji (https://imagej.net/Fiji). Axonal caliber (d) was calculated from the measured area (A) using the equation:

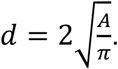

The inner tongue area was independently plotted as average per animal. Due to occurrence of phenotype at the inner tongue, the area including phenotypical *shiverer*-like membranes and the axon (A_n_) was subtracted using the following equation to obtain the corrected fiber caliber D_corr_:

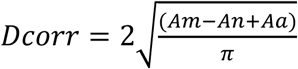

with A_m_: area compact myelin, A_n_: area non-compact myelin and axon, A_a_: axon area. The corrected g-ratio was then calculated as 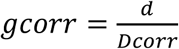

Analysis was performed on at least 5 FOV with at least 150 axons per animal on 3-4 animals.

### Immunoelectron microscopy

Immunogold labeling of cryosections prepared according to the Tokuyasu method was performed as previously described (Peters and Pierson, 2008, Weil et al., 2019). Optic nerves were dissected and immersion fixed in 4% PFA + 0.25% glutaraldehyde in 0.1 M phosphate buffer overnight and cryo-protected using 2.3 M sucrose in 0.1 M phosphate buffer mounted onto aluminum pins for cryo-sectioning and frozen in liquid nitrogen. Ultrathin 50-80 nm cryosections sections were cut with a 35° diamond knife, cryo-immuno 2,0 mm (Diatome, Biel, Switzerland) using a Leica UC6 ultramicrotome with a FC6 cryochamber (Leica, Vienna, Austria). Primary antibodies used were specific for PLP (A431, (Jung et al., 1996) and MBP (Custom made MBP antibody, this study). Protein A-gold conjugates were obtained from the Cell Microscopy Center, Department of Cell Biology, UMC Utrecht, The Netherlands (https://www.cellbiology-utrecht.nl/products.html). Sections were imaged using a LEO EM912 Omega transmission electron m)croscope (Carl Zeiss Microscopy, Oberkochen, Germany). For quantification of MBP density at least 4 sections with in total 300 μm^2^ per animal were quantified using a 2 μm grid to randomly select axons (n =3-4 animals, optic nerve, 26 weeks pti). Gold particle number and compact myelin area per sheath were counted for every randomly selected axon using Fiji (Schindelin et al., 2012) and Microscopy image browser (Belevich et al., 2016). Graphs display gold particles per μm^2^ compact myelin. Quantifications were performed blinded to the genotype and statistical analysis was performed using a two-tailed unpaired t-test in GraphPad prism 7.0 comparing control to iKO.

### Focused ion beam-scanning electron microscopy (FIB-SEM)

FIB-SEM was performed as described in (Steyer et al., 2019b, Weil et al., 2018). To visualize the emergence of membrane structures along a myelinated internode we imaged optic nerves 16 weeks and 26 weeks after tamoxifen induction. Optic nerves were either prepared by HPF and FS as described above or fixed for 24 h in 4% paraformaldehyde (PFA) and 2.5% glutaraldehyde in 0.1 M phosphate buffer. To achieve sufficient contrast for detection of backscattered electrons with the ESB detector chemically fixed optic nerves were processes using a modified protocol of the reduced osmium-thiocarbohydrazide-osmium (rOTO) method (Deerinck et al., 2010) as described previously (Erwig et al., 2019b). Nerves were transferred into embedding molds filled with Durcupan and polymerized at 60°C for 48h as previously described. Samples for FIB-SEM imaging of high-pressure frozen optic nerve were prepared as described above and embedded in Durcupan (Sigma -Aldrich) instead of Epon. Samples in blocks were then trimmed using a 90° diamond trimming knife (Diatome AG, Biel, Switzerland) and mounted on a SEM stub (Science Services GmbH, Pin 12.7 mm x 3.1 mm) using a silver filled epoxy resin (Epoxy Conductive Adhesive, EPO-TEK EE 129–4; EMS) and polymerized at 60° overnight.

Optic nerves used for quantification of myelin coverage were minimal embedded in Durcupan as described (Steyer et al., 2019a, Schieber et al., 2017) and polymerized at 60°C for 48h. Polymerized nerves were then also mounted on a SEM stub (Science Services GmbH, Pin 12.7 mm x 3.1 mm) using a silver filled epoxy resin (Epoxy Conductive Adhesive, EPO-TEK EE 129–4; EMS) and polymerized at 60° overnight. All samples were coated with a 10 nm platinum layer using a sputter coater EM ACE600 (Leica) at 35 mA current. Samples were placed into the Crossbeam 540 focused ion beam-scanning electron microscope (Carl Zeiss Microscopy GmbH). To protect the sample surface, a 300-500 nm platinum or carbon layer was deposited on top of the region of interest. Atlas 3D (Atlas 5.1, Fibics, Canada) software was used for milling and collection of 3D data. Initial milling was performed with a 15 nA current followed by a 7 nA current to polish the surface. Imaging was performed at 1.5 kV using an ESB detector (450 V ESB grid, pixel size x/y 5 nm) in a continuous mill-and-acquire mode using 700 pA for the milling (z-step 50 nm).

Images were aligned using the ImageJ plugin TrackEM2 (Cardona et al., 2012) followed by postprocessing in Fiji: Images were cropped, inverted and blurred (Gaussian blur, sigma 2) to suppress noise. Stacks were manually segmented using IMOD (Kremer et al., 1996). Quantification of phenotypes in stacks of optic nerves 16W and 26W post tamoxifen were performed manually using Microscopy image browser (Belevich et al., 2016). Myelin coverage and number of myelin spheres was measured in FIB-SEM stacks in n=1 animals and stacks at 26 weeks pti with at least 100 axons and at 16 weeks pti in n=2 animals with at least 90 axons per animal.

### Nanoscale Secondary Ion Mass Spectrometry (Nano-SIMS) imaging

Semithin sections of Epon-embedded spinal cord samples were collected on finder grids (formvar and carbon-coated 200 square mesh copper grids, FCF200-1-Cu, Science Services, Munich, Germany) and regions of interest were mapped by taking images at increasing magnification by TEM. NanoSIMS imaging was performed as previously described (Kabatas et al., 2019, Saka et al., 2014) by a nanoSIMS 50L (CAMECA, Gennevilliers Cedex, France) with an 8 kV Cesium primary source. To detect the presence of ^12^C and ^13^C, the signals of ^12^C^14^N^-^ and ^13^C^14^N^-^ were measured. To reach the steady state of ionization, the samples were first implanted with a primary current of ∼15 pA. A current of ∼ 0.5-1 pA was applied during the imaging. Entrance slit and aperture slit were selected to obtain sufficient mass resolving power for a good separation of ^13^C^14^N^-^ peak from the interference peaks ^12^C^15^N^-^. The images were obtained with the raster size between 10⨯10 μm and 20×20 μm and 256×256 pixels, or the size lager than 20×20 μm and 512×512 pixels. Image exportation, layer addition and drift correction, were performed by the OpenMIMS plugin from Fiji (Schindelin et al., 2012) and self-written Matlab (the Mathworks Inc, Natick, MA) scripts were used for the analysis and correlation of EM and SIMS images. To analyze the ^13^C enrichment in different regions, the TEM images were first aligned with the nanoSIMS images. ROIs were then manually drawn by an expert on the TEM image. From the ROIs the isotopic ratio of ^13^C^14^N^-^ / ^12^C^14^N^-^ was extracted to evaluate the ^13^C enrichment. The ratio was calculated for each pixel in the ROI, and then the average across all pixels in the ROI was calculated and presented. We employed T-tests to determine if the ^13^C enrichment was significantly different between regions. The standard for ^13^C/^12^C natural ratio (0.0112) was set as 0% enrichment.

## QUANTIFICATION AND STATISTICAL ANALYSIS

The group size (number of animals = n) and the statistical test used are indicated in the respective figure legend. For calculation of SD and testing for significance MSExcel and Graphpad Prism were used.

### Analysis of proteomic data

For label-free protein quantification, the freely available software ISOQuant (www.isoquant.net) was used for post-identification analysis including retention time alignment, exact mass and retention time (EMRT) and ion mobility clustering, peak intensity normalization, isoform/homology filtering and calculation of absolute in-sample amounts for each detected protein (Kuharev et al., 2015, Distler et al., 2014, Distler et al., 2016). Only peptides with a minimum length of seven amino acids that were identified with scores above or equal to 5.5 in at least two runs were considered. FDR for both peptides and proteins was set to 1% threshold and only proteins reported by at least two peptides (one of which unique) were quantified using the TOP3 method (Silva et al., 2006). The parts per million (ppm) abundance values (i.e. the relative amount (w/w) of each protein in respect to the sum over all detected proteins) were log2-transformed and normalized by subtraction of the median derived from all data points for the given protein. As described in detail recently (Ambrozkiewicz et al., 2018), significant changes in protein abundance were detected by moderated t-statistics across all technical replicates using an empirical Bayes approach and false discovery (FDR)-based correction for multiple comparisons (Kammers et al., 2015), realized in the Bioconductor R packages limma and q-value. The relative abundance of a protein was accepted as altered for q-values <0.05. To detect changes in normalized protein abundance over the course of MBP deficiency, we used limma to analyze the difference of differences with the interaction term (iKO40w-Ctrl40W)-(iKO8w-Ctrl8W) according to the limma User’s Guide (https://bioconductor.org/packages/release/bioc/vignettes/limma/inst/doc/usersguide.pdf). The exact q-values are reported in Supplementary Table 1.

## DATA AND CODE AVAILABILITY

The datasets generated and/or analyzed during the current study are available from the corresponding author upon reasonable request. Upon acceptance we plan to deposit the original data sets in this accessible archive: https://www.ebi.ac.uk/pdbe/emdb/empiar/.

The Matlab (the Mathworks Inc, Natick, MA) scripts used for analysis of NanoSIMS data can be obtained from Silvio Rizzoli.

### Key Resources Table

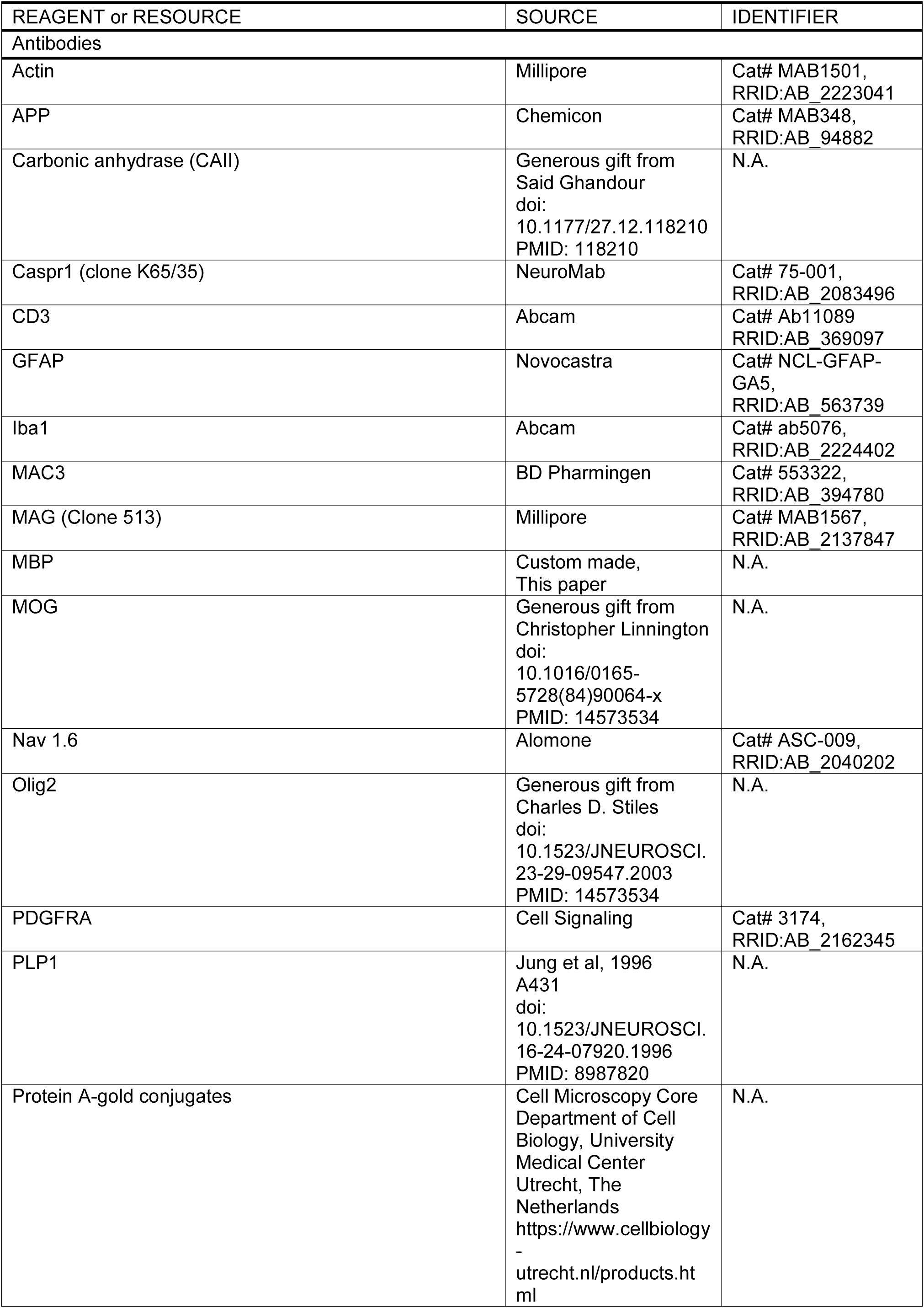
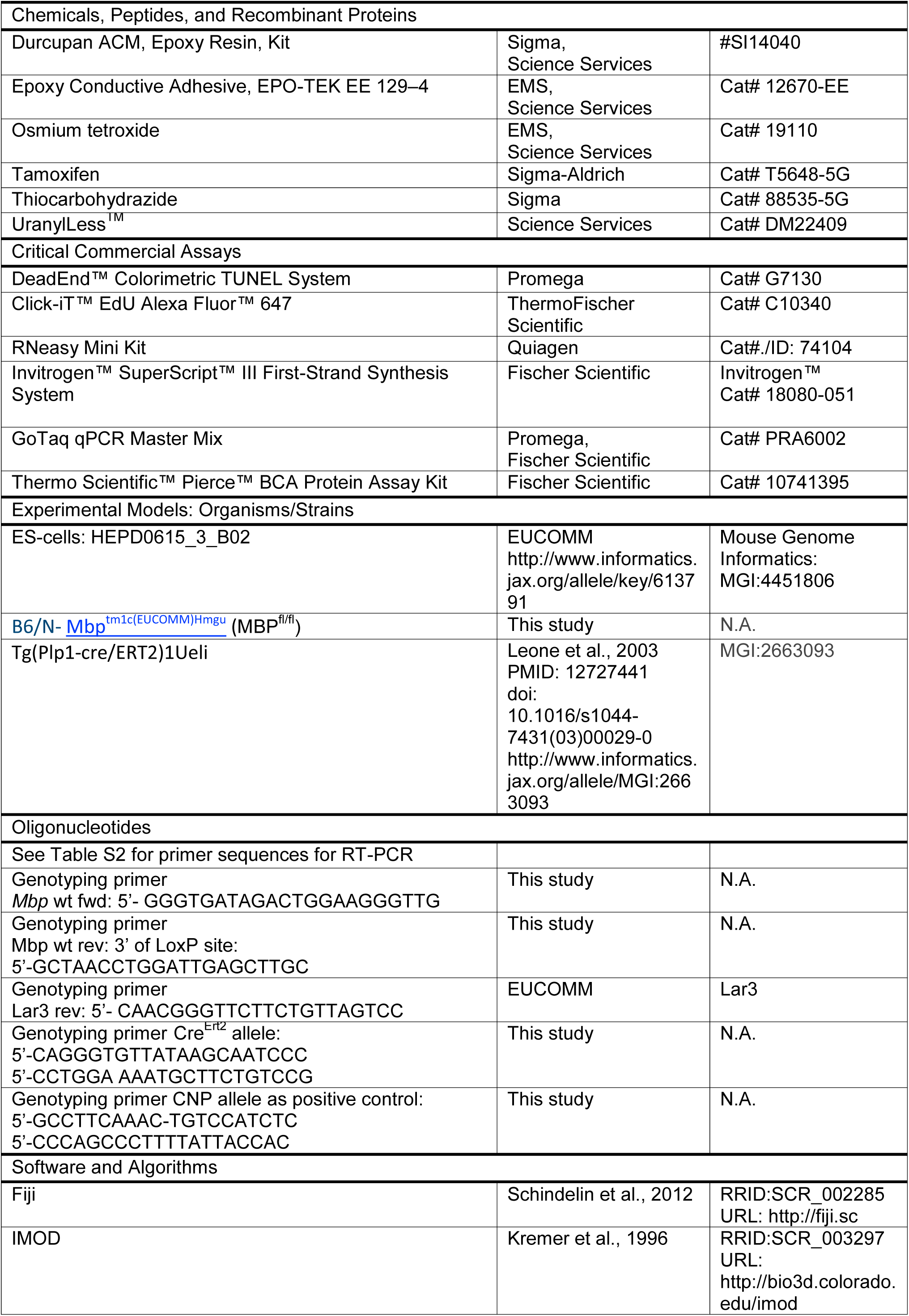
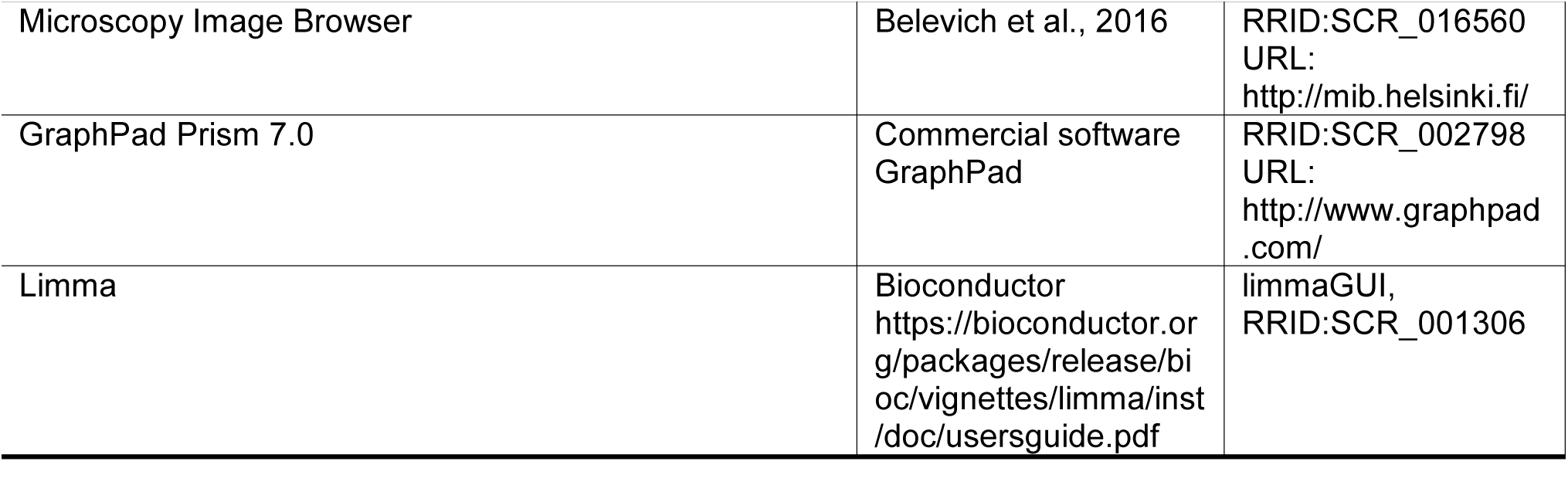

### Supplementary movies S1-S4

**Movie S1 related to Figure 5: Internode shortening**

FIB-SEM data stack and 3D reconstruction of a myelinated axon showing the pathological phenotype: The residual internode (in red) is short, shiverer-like membranes processes occur at the juxtaparanode and also emerge at the paranode and bend away from the axon (indicated in yellow). The image stack was recorded with a voxel size of 5 nm ⨯ 5 nm ⨯ 50 nm; the movie consists of 747 images cropped from the original data stack. The optic nerve was fixed chemically and embedded using the reduced osmium-thiocarbohydrazide-osmium (rOTO) method (Deerinck et al., 2010). The myelinated axon was reconstructed using IMOD (Kremer et al., 1996).

**Movie S2 related to Figure 6: Myelin tubulation at the juxtaparanode**

FIB-SEM data stack with indicated phenotype of myelin tubulation occurring at the juxtaparanode. The data stack was recorded with a voxel size of 5 nm x 5 nm x 50 nm. The movie consists of 318 images which were colored using IMOD. The optic nerve sample was prepared by high-pressure freezing and freeze substitution (HPF/FS) at 16 weeks pti.

**Movie S3: Axonal wrapping in shiverer optic nerve**

FIB-SEM data stack of an optic nerve sample of a shiverer mouse at 8 weeks of age. Coloration indicates an axon and the associated shiverer membrane wrapping. The data stack was recorded with a voxel size of 5 nm ⨯ 5 nm ⨯ 50 nm. The sample was prepared by chemical fixation and rOTO embedding, coloration was performed in IMOD.

**Movie S4 related to Figure 7: Myelin thinning and formation of myelinoid bodies**

FIB-SEM data stack showing a myelinated axon 26 weeks pti. Substantial myelin thinning and the formation of myelinoid bodies are visible. The sample was prepared by chemical fixation and rOTO embedding. Outline of myelin (in yellow) and axon (in red) and myelinoid bodies in different colors. The images were recorded with a voxel size of with 5 nm x 5 nm x 50nm, the movie consists of 404 images cropped from the original data stack. Coloration was performed in IMOD.

