## Supplementary Figures_Meschkat et al for "White matter integrity requires continuous myelin synthesis at the inner tongue"

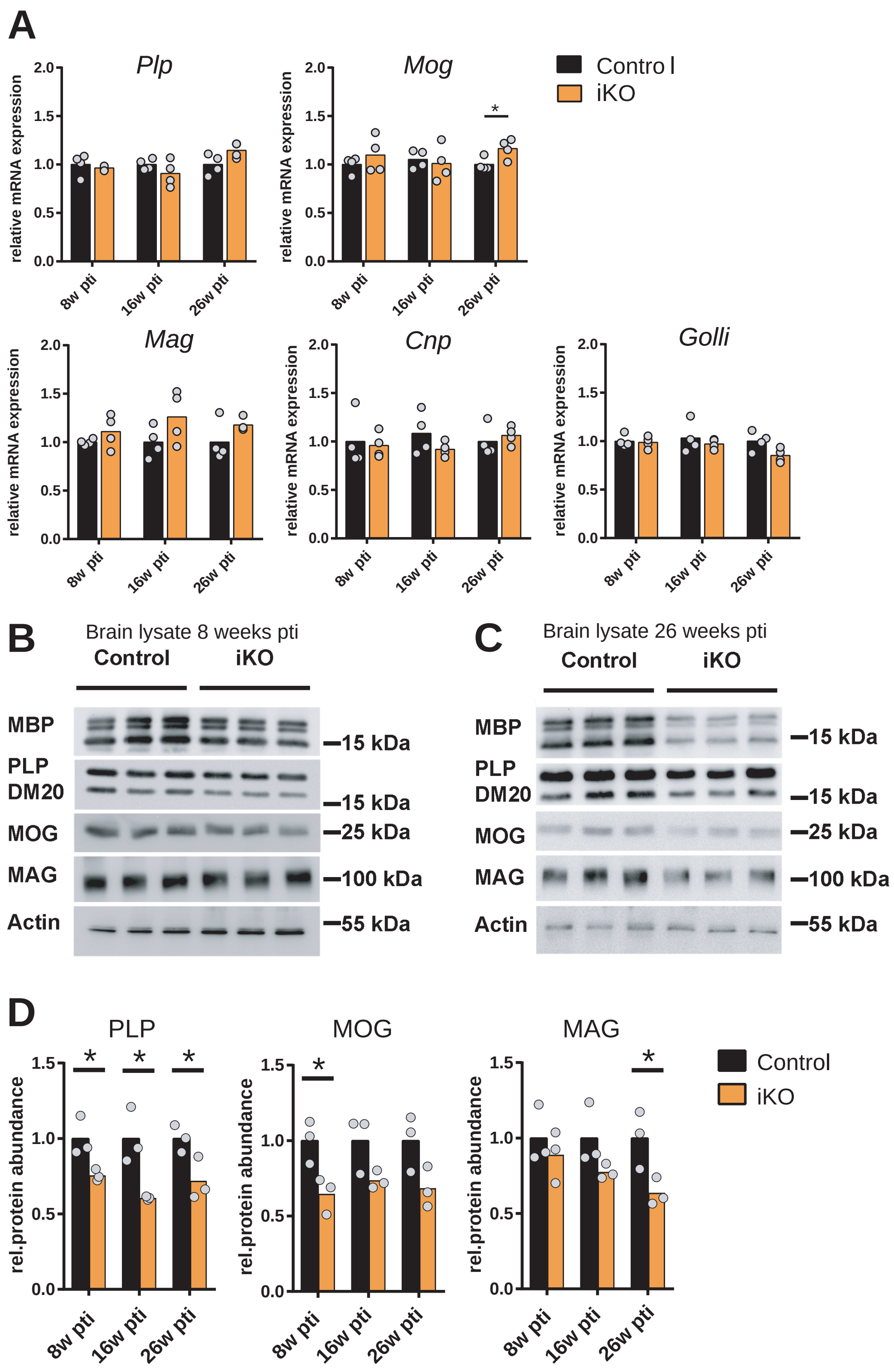


**Figure S1 (related to Fig 1D-F): Quantification of mRNA abundance of myelin genes and myelin proteins by immunoblotting at indicated time points after ablation of *Mbp***

**(A)** qRT-PCR of brain lysate showed that mRNA levels of indicated myelin genes are unchanged after ablation of *Mbp*. (Statistics: two-tailed unpaired t-test $p < .05$ (*), $p < .01$ (**), $p < .001$ (***), N=3-4 animals. **(B)** Immunoblot of brain lysate 8 weeks pti and **(C)** 26 weeks pti and quantitation in **(D)**. The protein abundance of PLP is reduced, for MOG and MAG trends towards decreased abundance is detectable: PLP 8 weeks pti (74%), 16 weeks pti (60%), 26 weeks pti (71%); MOG 8 weeks pti (64%), 16 weeks pti (73%), 26 weeks pti (68%); MAG 8 weeks pti (88%), 16 weeks pti (73%), 26 weeks pti (63%) (two-tailed unpaired t-test $p < .05$ (*)). For *Mbp* mRA abundance and MBP protein levels in brain lysate see Fig 1D and E and Fig. 1F and G respectively.

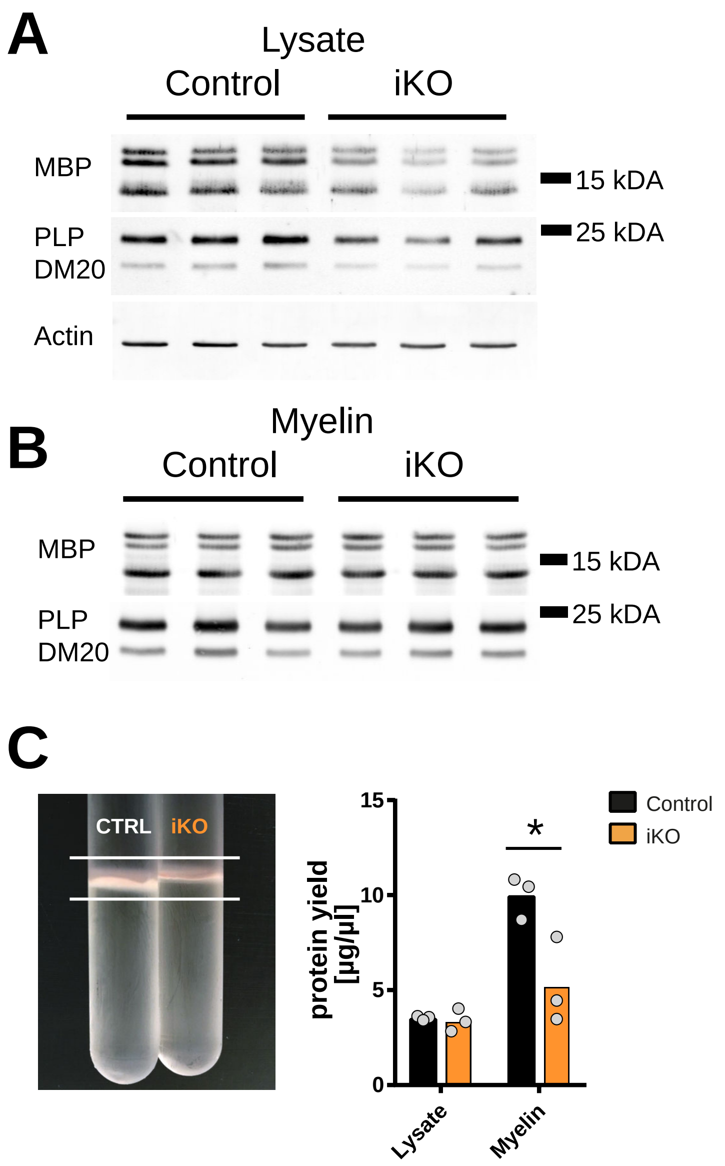


**Figure S2 (related to Fig. 1G): Abundance of MBP in biochemically purified myelin.**
**(A)** Immunoblot of MBP and PLP in Lysate and **(B)** myelin biochemically purified from brains (n=3) reveals a reduction on lysate level but no changes in MBP and PLP protein abundance in the myelin fraction. **(C)** Isolation of a myelin enriched fraction shows a visibly reduced yield of myelin membrane and protein from iKO brains 26 weeks pti, Bradford protein assay of lysate and myelin (two-tailed unpaired t-test $p < .05$ (*)).

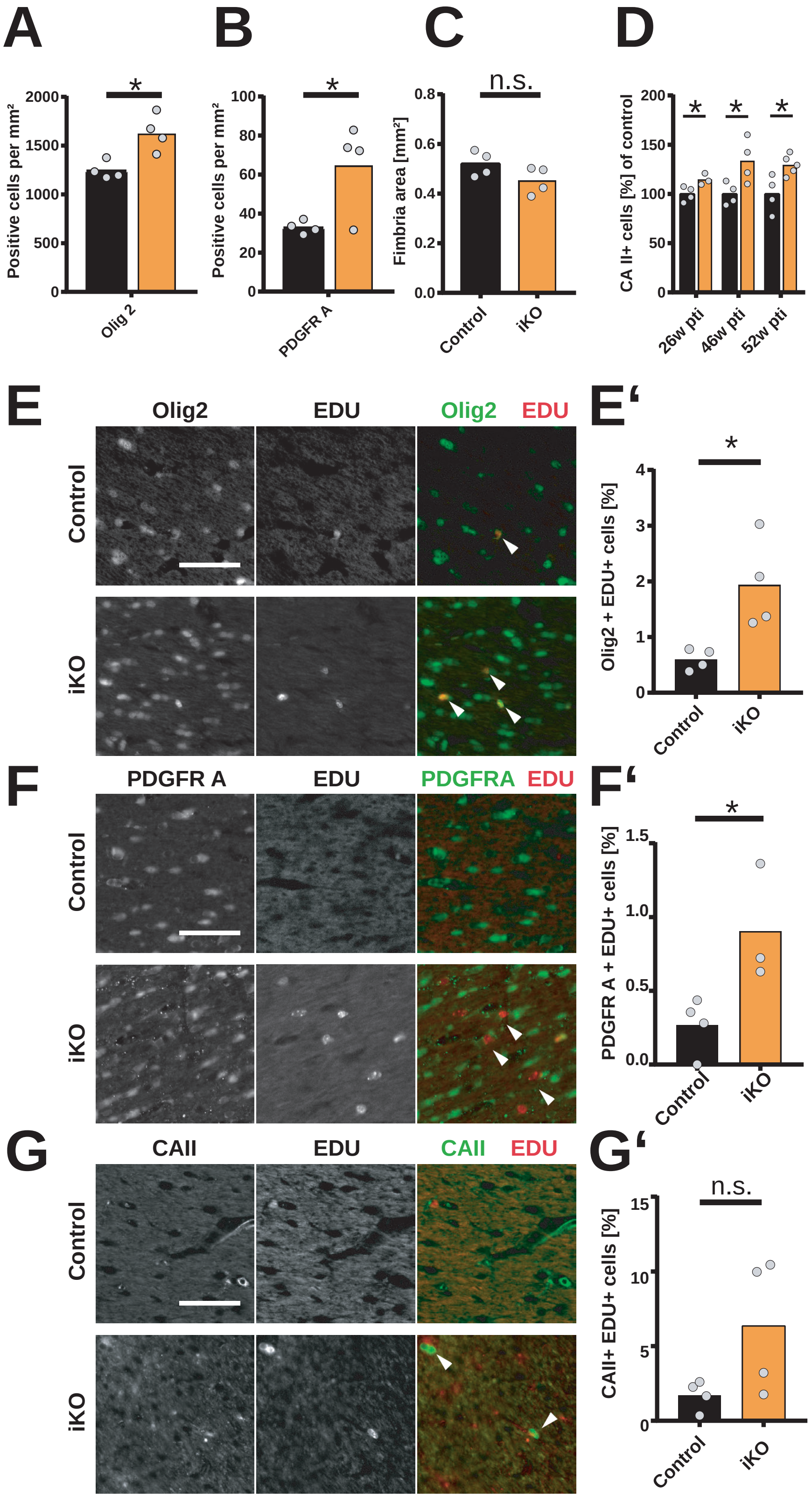


**Figure S3 (related to Fig. 1): Oligodendrocyte numbers in the fimbria are increased after *Mbp* ablation.** **(A)** The densities of Olig2$^{+}$ and **(B)** PDGFRA$^{+}$ cells were increased 46 weeks pti while the fimbria area was not significantly decreased **(C)**. Quantification of CAII$^{+}$ cells at 26, 46 and 52 weeks pti revealed a consistent increase in CAII$^{+}$ cells **(D)**. Mice at 40 weeks pti were treated with 0.2 mg/ml EdU in drinking water for three weeks followed by a three weeks chase period to allow cell maturation of labeled cells. Immunohistochemistry of cells positive for EDU$^{+}$ and oligodendrocyte linage marker Olig2 **(E)**, the OPC marker PDGFRA **(F)** and the mature oligodendrocyte marker CAII (**G**). Quantification showed a significant increase in EDU$^{+}$ Olig 2$^{+}$ and EDU$^{+}$ PDGFR A$^{+}$ double positive cells **(E', F')**. **(E', F', G')** % of EDU positive cells from Olig 2, PDGFRA or CAII positive cells per FOV, N= 3-4 animals, two-tailed unpaired *t*-test; $p < .05$ (*), $p < .01$ (**), $p < .001$ (***). Scale bars: 50 µm.

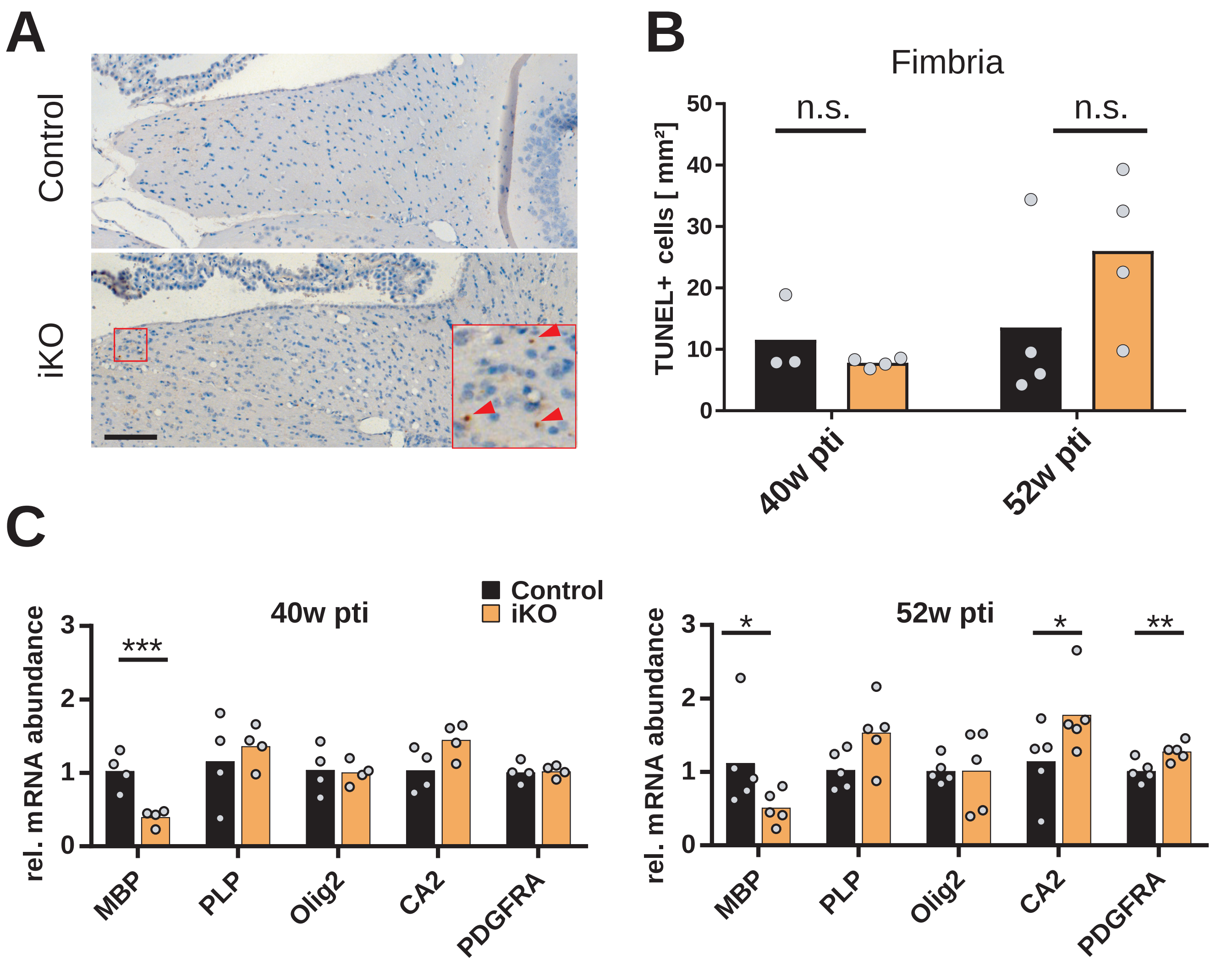


**Figure S4 (related to Fig. 1): Survival of recombined oligodendrocytes.**
**(A)** TUNEL cell death assay to test for increased cell death in the fimbria of iKO animals showed no significant increase in TUNEL-positive cells at 40 and 52 weeks pti depicted in **(B)**. Chromogenic labeling, N= 3-4 animals, 2 fimbria per animal, two-tailed unpaired *t*-test, scale bar 100 µm. **(C)** qRT-PCR analysis of corpus callosum at 40 and 52 weeks pti. Expression of *Mbp* mRNA is reduced to 50% of control while the expression of *Plp* and *Olig 2* is unchanged. The OPC marker *PDGFRA* and the mature oligodendrocyte marker *CAII* show increased expression at 52 weeks pti (n= 4-5 animals, two-tailed unpaired t-test, p < .05 (*), p < .01 (**), p < .001 (***).

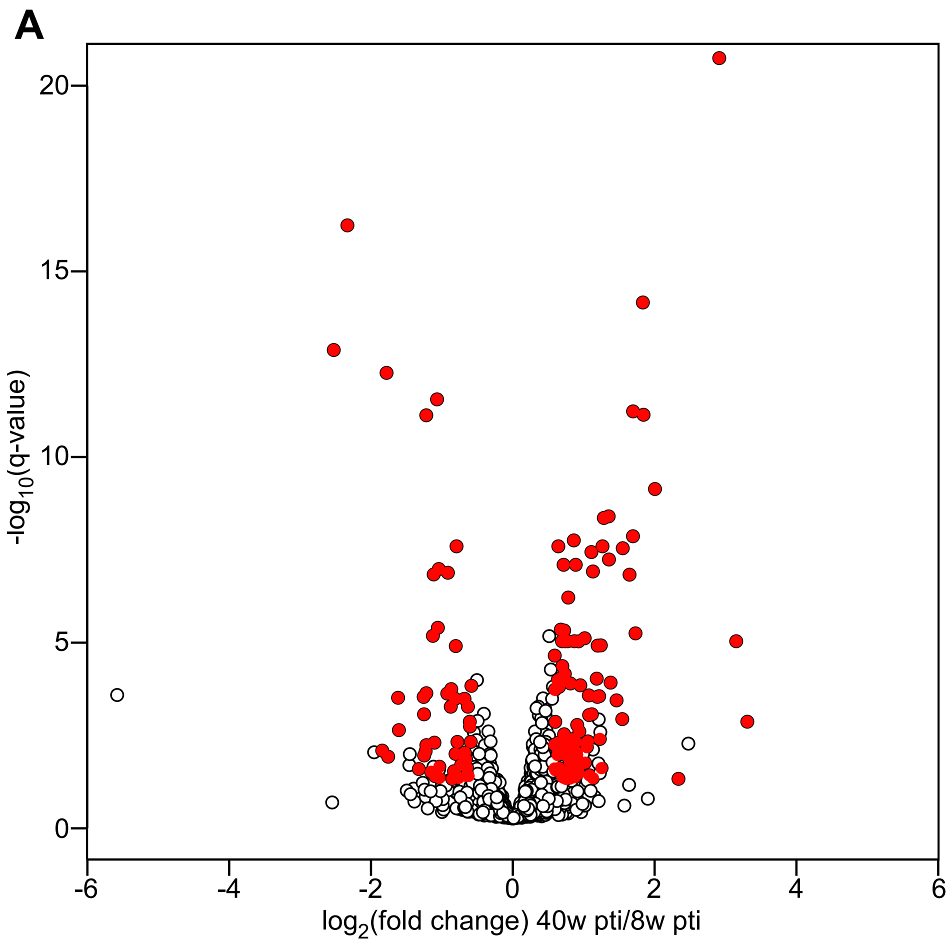


**Figure S5 (related to Fig. 1): Proteome alterations in the late stage of MBP deficiency.**
**(A)** Volcano plot visualizing differences in normalized protein abundance between early (iKO/Ctrl 8w) and late (iKO/Ctrl 40w) time points after MBP deletion. From the 1863 proteins in the entire dataset (open black circles), 149 proteins remained after stringent filter criteria: log2 fold-change (FC) >|0.58|, q-value < 0.05, min. 6 out of 8 values detected per group (filled red circles, see also Supplementary Table 1). As no imputation of missing values was performed, proteins exclusive for iKO or Ctrl are not part of the analysis and were considered separately by manual analysis. **(B)** Clustering of the normalized abundance of the 149 proteins derived from the volcano plot in **(A)**. The heatmap guided the selection of categories and their protein representatives for the visualization of pathological events in Fig. 1I-K. Selected proteins are shown in red, with an enlarged protein acronym (nomenclature as used in Fig. 1I-K) shown adjacent to the protein ID column of the heatmap

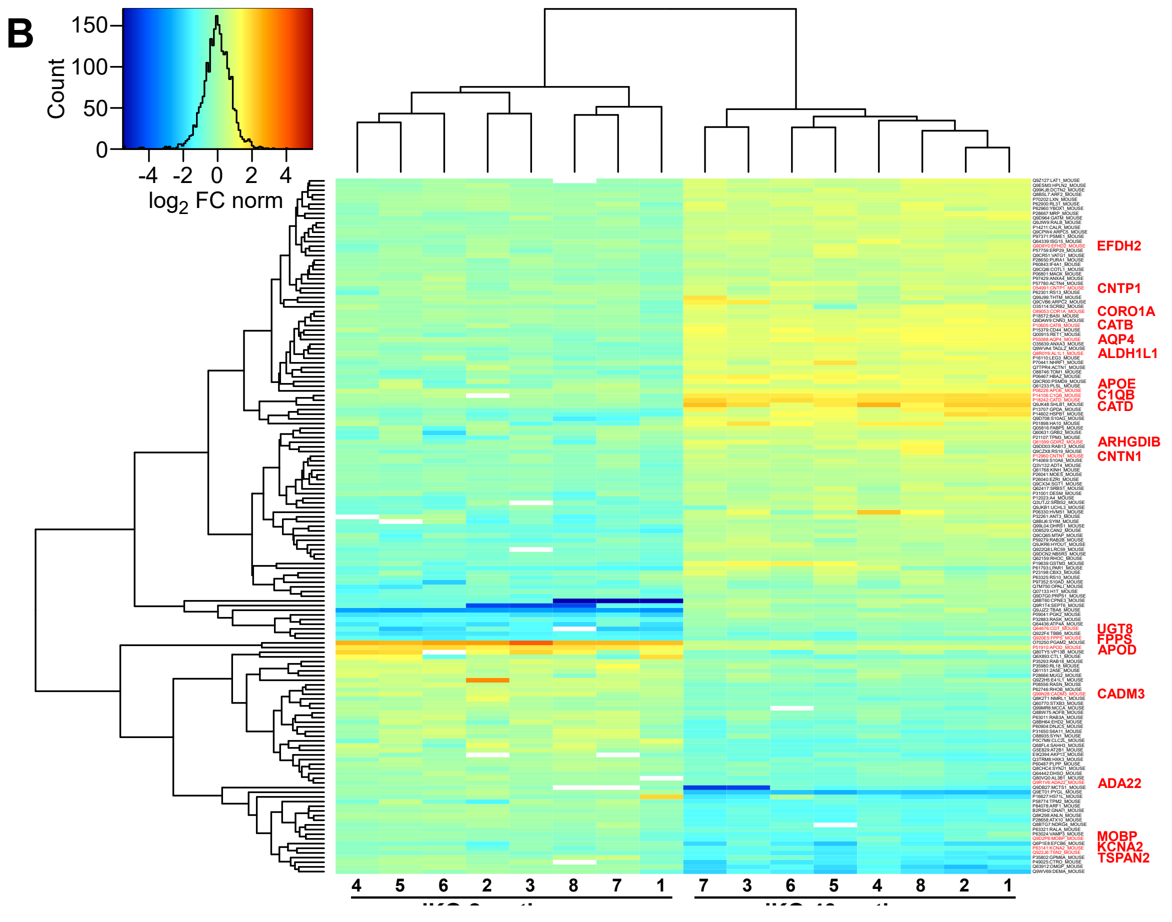

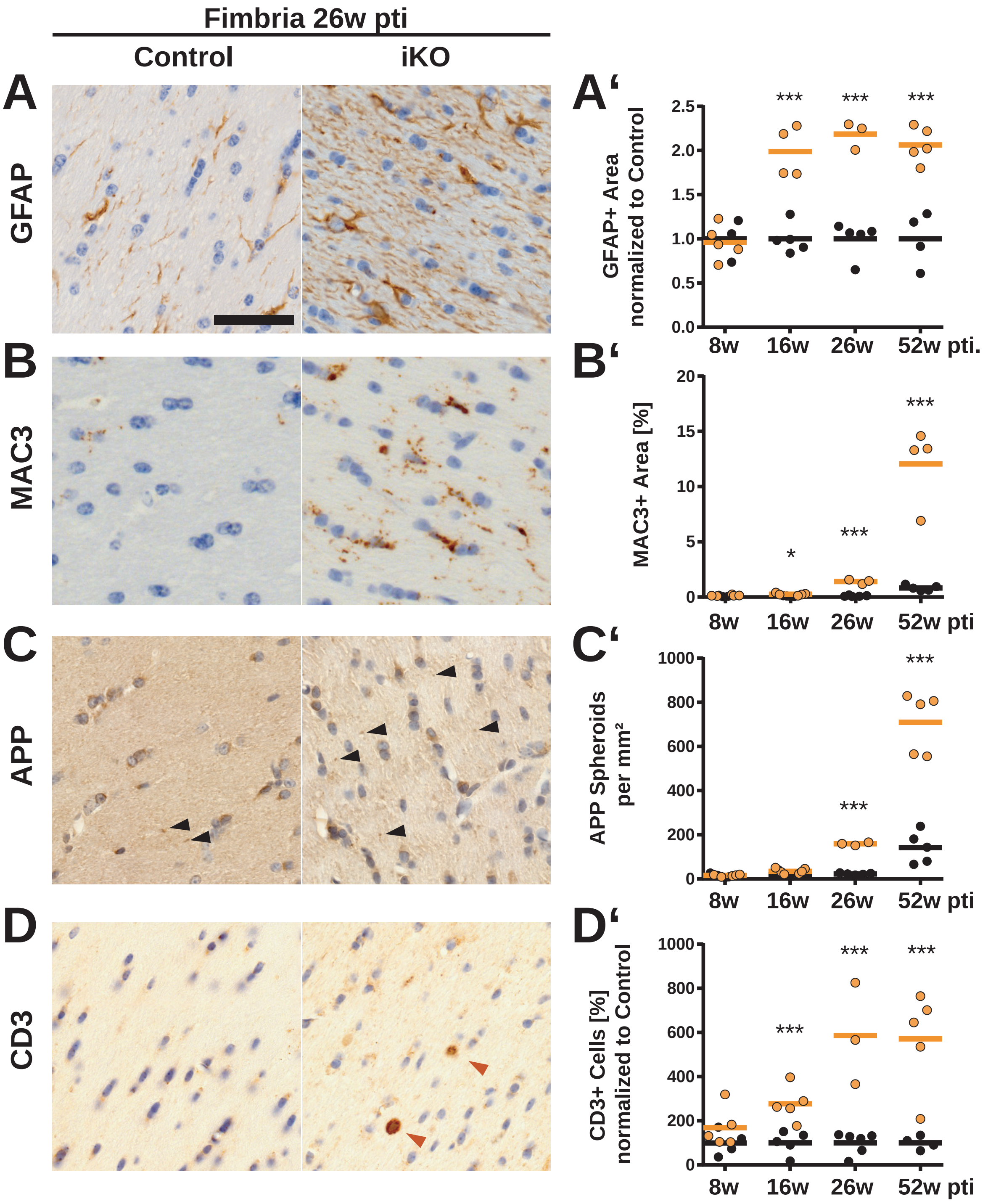


**Figure S6 (related to Fig. 1): Slow development of inflammation and neuropathology in the fimbria.**
**(A-D)** Immunhistochemical staining of GFAP, MAC3, APP and CD3 in the fimbria 26 weeks pti and in **(A'-D')** quantification of indicated time points. An increase in GFAP$^{+}$ area **(A)** indicates astrogliosis beginning between 8 and 16 weeks pti **(A')**. The increase of MAC3$^{+}$ area **(B)** indicates microglial activity beginning between 8 and 16 weeks pti and increasing until 52 weeks pti **(B')**. Axonal pathology observed by labeling of APP spheroids **(C)** appears late at 26 weeks pti and increases until 52 weeks pti **(C')**. Increased numbers of T-cells **(D)** can be observed already at 16 weeks pti indicating immune response (cells/mm$^2$ normalized to control) **(D')**. N=3-5 animals, 2 fimbria (GFAP, MAC3, APP) or 6 (CD3) fimbria per animal, two-tailed unpaired *t*-test; $p < .05$ (*), $p < .01$ (**), $p < .001$ (***). Scale bars: 50 µm

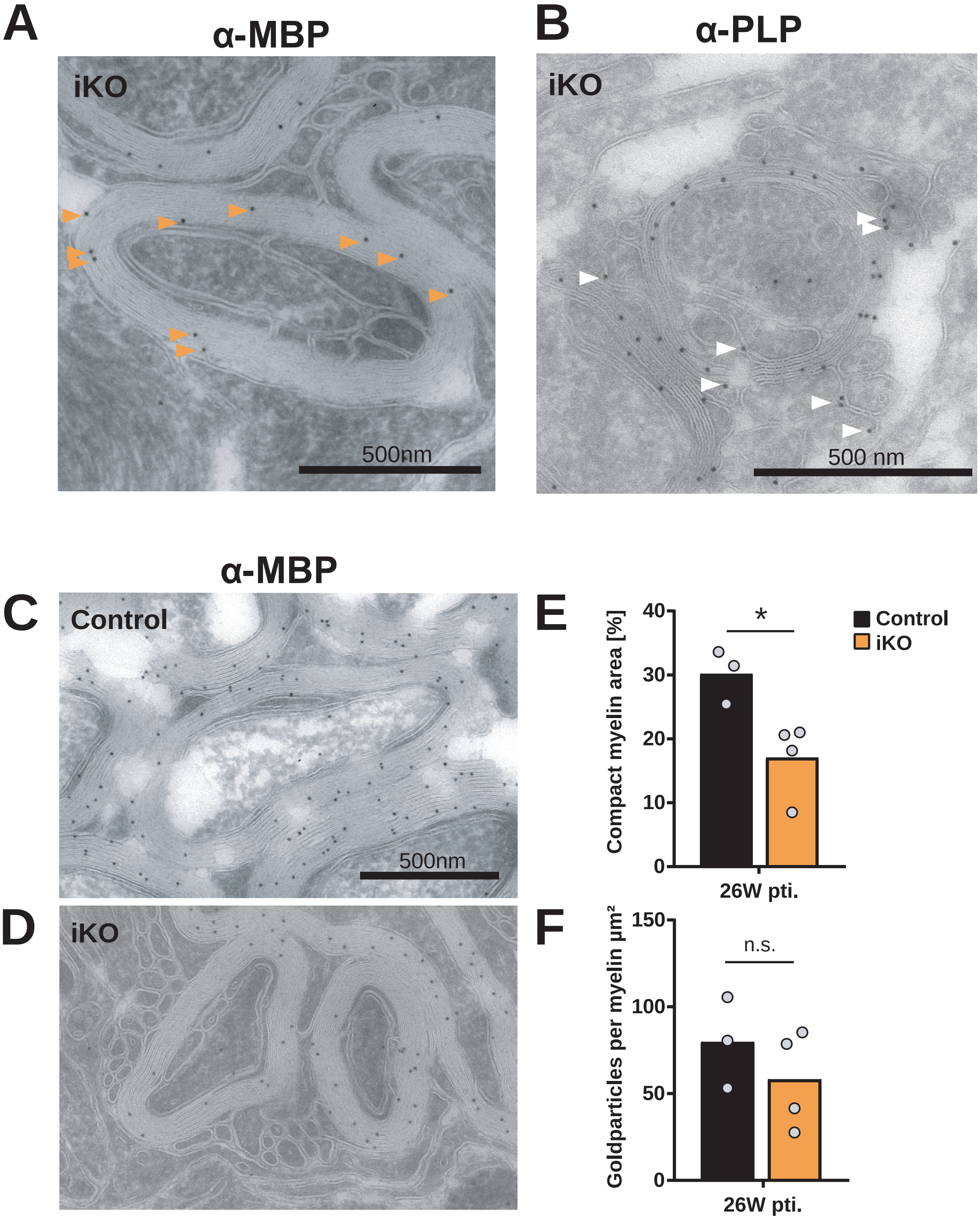


**Figure S7 (related to Fig. 3): Membrane tubules are oligodendrocytic membranes that are devoid of MBP.**
**(A-B)** Membrane tubules found at the inner tongue and adjacent to axons are MBP negative **(A)** but show PLP-labeling **(B)** indicating an oligodendrocytic origin. Optic nerve 26 weeks pti, 10 nm protein-A gold particles bound to primary antibody against PLP and MBP. **(C-D)** The labeling density of MBP in control myelin **(C)** is comparable to iKO myelin **(D)** suggesting little reduction in MBP content within the compact myelin of iKOs. **(F)** The section area occupied by myelin is decreased after *Mbp* iKO **(E)**. 10 nm protein-A-gold particles. At least 4 sections with in total 300 µm$^2$ per animal were quantified using a 2 µm grid to randomly select axons, two-tailed unpaired t-test; $p < .05$ (*). Scale bar: 500 nm

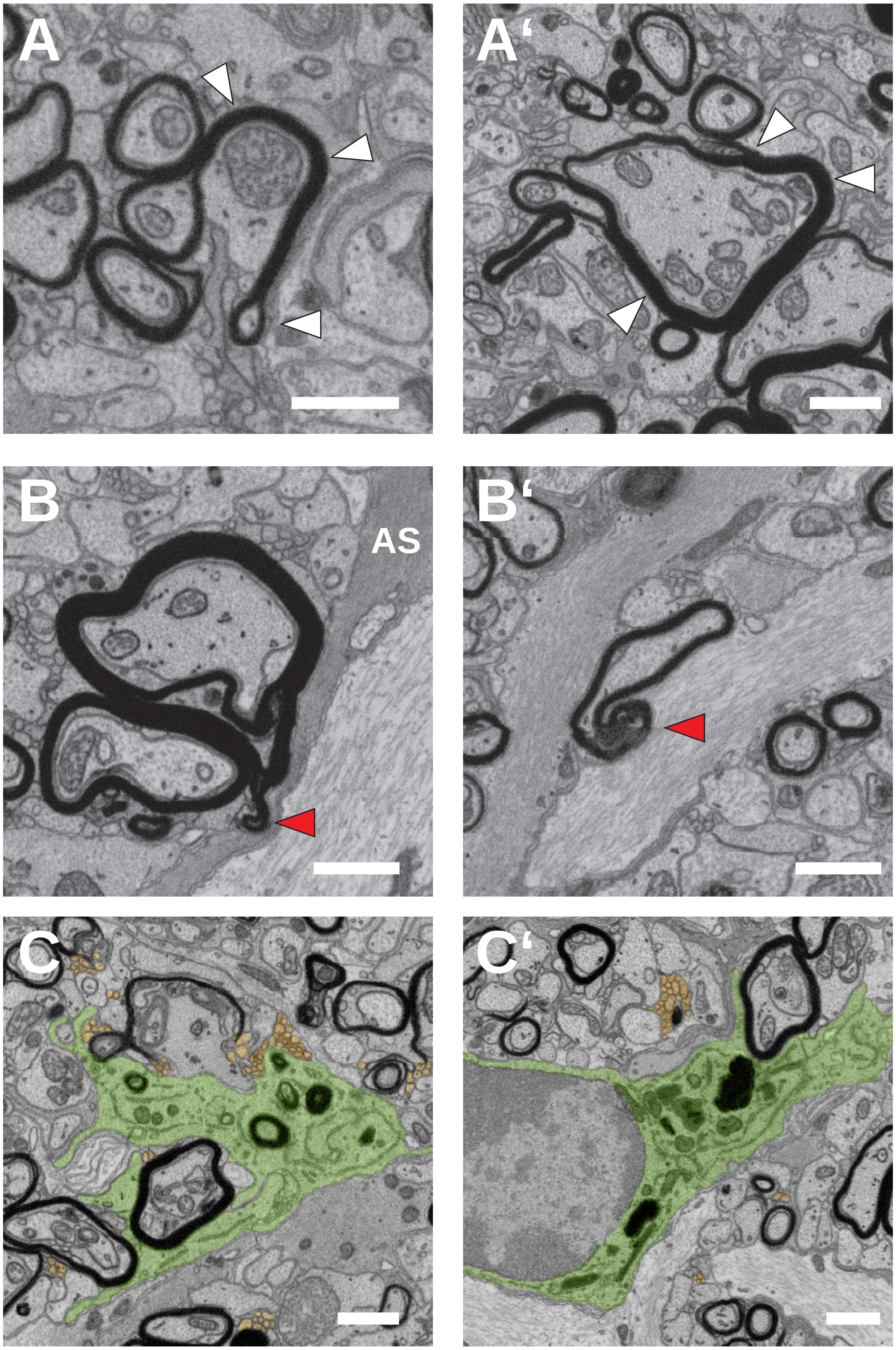


**Figure S8 (related to Fig. 7): Myelin outfoldings engulfed by astrocytes and myelin debris incorporated by microglia** **(A, A')** Myelin outfoldings (white arrow head) are visible in the optic nerve 26 weeks pti. **(B-B')** Astrocytes (AS) are found in direct contact to myelin protrusions (red arrow head). **(C-C')** Microglia (false colored in green) are found close to demyelinating axons (asterisk) and non-compact myelin membrane (false colored in orange). Presence of dark degradative compartments containing myelin debris indicate microglial uptake of myelin. Images were extracted from a FIB-SEM image stack of optic nerve 26 weeks pti. Scale bars: 1 µm.
